# A late-stage assembly checkpoint of the human mitochondrial ribosome large subunit

**DOI:** 10.1101/2021.06.09.447755

**Authors:** Pedro Rebelo-Guiomar, Simone Pellegrino, Kyle C. Dent, Aldema Sas-Chen, Leonor Miller-Fleming, Caterina Garone, Lindsey Van Haute, Jack F. Rogan, Adam Dinan, Andrew E. Firth, Byron Andrews, Alexander J. Whitworth, Schraga Schwartz, Alan J. Warren, Michal Minczuk

## Abstract

The epitranscriptome plays a key regulatory role in cellular processes in health and disease, including ribosome biogenesis. Here, analysis of the human mitochondrial transcriptome shows that 2’-*O*-methylation is limited to residues of the mitoribosomal large subunit (mtLSU) 16S mt-rRNA, modified by MRM1, MRM2, and MRM3. Ablation of MRM2 leads to a severe impairment of the oxidative phosphorylation system, caused by defective mitochondrial translation and accumulation of mtLSU assembly intermediates. Structures of these particles (2.58 Å) present disordered RNA domains, partial occupancy of bL36m and bound MALSU1:L0R8F8:mtACP anti-association module. Additionally, we present five mtLSU assembly states with different intersubunit interface configurations. Complementation studies demonstrate that the methyltransferase activity of MRM2 is dispensable for mitoribosome biogenesis. The *Drosophila melanogaster* orthologue, *DmMRM2*, is an essential gene, with its knock-down leading to developmental arrest. This work identifies a key late-stage quality control step during mtLSU assembly, ultimately contributing to the maintenance of mitochondrial homeostasis.

## INTRODUCTION

The human mitochondrial genome is encoded in multiple copies of ∼16.6 kb circular double-stranded DNA molecules (mtDNA) present in mitochondrial nucleoids in the mitochondrial matrix. Expression of this genome entails several, highly regulated processes, with newly synthesised transcripts being cleaved, chemically modified, polyadenylated and further matured in neighbouring structures known as mitochondrial RNA granules (MRGs). The assembly of mitochondrial ribosomes also takes place in MRGs, with mitoribosomal proteins (MRPs) and biogenesis factors engaging in a complex process, which has not yet been fully characterised (Bogenhagen, Martin and Koller, 2014; Antonicka and Shoubridge, 2015; Jourdain et al., 2016).

Similar to other systems, the mitochondrial ribosome is composed of a small (mtSSU) and a large (mtLSU) subunit, with their core rRNAs, 12S and 16S mitochondrial (mt-) rRNAs, respectively, surrounded by MRPs (30 for the mtSSU and 52 for the mtLSU). While the RNA components of the mitoribosome are mitochondrially-encoded, all MRPs and assembly factors are encoded in the nuclear genome, thus requiring coordination between two genomes for the assembly of these macromolecular complexes. The mammalian mitochondrial ribosome is endowed with a number of specific features. While RNA makes up most of the composition of bacterial and cytosolic eukaryotic ribosomes, mammalian mitochondrial ribosomes present a more elaborate protein shell, which aids coping with the oxidative microenvironment. Almost half of these MRPs are evolutionarily exclusive to mitochondrial ribosomes, some of which were repurposed and accreted during reductive genome evolution (Brown et al., 2014; Petrov et al., 2019). In addition, several mitochondrial ribosomal proteins that share homology to other translation systems present mitochondria-specific extensions, structurally compensating truncated rRNA segments (Sharma et al., 2003; Brown et al., 2014; Amunts et al., 2015; Petrov et al., 2019). The peptide exit tunnel is lined with hydrophobic residues, which stabilise the highly hydrophobic nascent mitochondrial peptides (Brown et al., 2014). Furthermore, instead of 5S rRNA, structurally similar mtDNA-encoded tRNAs occupy an equivalent region in the central protuberance of the mtLSU. Depending on the organism and availability, mt-tRNA^Val^ or mt-tRNA^Phe^ are incorporated, most likely due to their genomic proximity to mt-rRNA genes and consequent near stoichiometric presence of their transcripts (Brown et al., 2014; Rorbach et al., 2016).

As other RNA classes, mt-rRNAs contain modified ribonucleotides which are post-transcriptionally introduced by a set of enzymes. Unlike the bacterial and eukaryotic cytoplasmic counterparts which rRNAs carry tens of post-transcriptional modifications, there are only 10 modified residues in mammalian mt-rRNAs, all clustering in functionally relevant regions of the mitoribosome, including the A- and P-loops of the peptidyl transferase centre (PTC) in the 16S mt-rRNA, or the decoding centre in the 12S mt-rRNA (Rebelo-Guiomar et al., 2019). Methylation of the 2’-hydroxyl group of ribonucleotides (2’-*O*-methylation) has been detected in G1145, U1369 and G1370 of 16S mt-rRNA (human mtDNA numbering) and the involved enzymes identified as MRM1, MRM2 and MRM3, respectively (Lee et al., 2013; Lee and Bogenhagen, 2014; Rorbach et al., 2014). However, the role of the modifications and the enzymes introducing them has remained elusive and lacking molecular and mechanistic insight. Furthermore, mutations in *MRM2* have been implicated in human pathogenesis, in an ever-growing group of patients affected by mitochondrial diseases, for which pathogenesis mechanisms are not yet completely understood with molecular detail (Garone et al., 2017).

With this, we set out to investigate the role of MRM2 in the biogenesis and function of human mitochondrial ribosomes. We show that the presence of MRM2, but not its methyltransferase activity, is essential for mtLSU biogenesis, constituting a quality control point in the late stages of this process. In the absence of MRM2, a severe mitochondrial dysfunction phenotype with underlying defects in mitochondrial translation is observed, with the mtLSU being trapped in immature assembly states where domains IV and V of 16S mt-rRNA are unstructured, and an anti-association module is present, preventing premature engagement of these immature particles in translation. Finally, using a *Drosophila melanogaster* model, we show that MRM2 is essential for development and homeostasis of neuronal and muscular tissues, often affected in mitochondrial diseases.

## RESULTS

### 2’-*O*-Methylation in the human mitochondrial transcriptome

To investigate the presence of nucleotides with 2’-*O*-methylated riboses in mitochondrial RNAs, we performed a transcriptome-wide profiling in cells lacking three known 2’-*O*-methlyltransferases, MRM1, MRM2 and MRM3 (**Figure 1** and **Figure S1**; Lee et al., 2013; Lee and Bogenhagen, 2014; Rorbach et al., 2014). To this end, we used a combination of two high throughput sequencing-based methods. First, an adaptation of RiboMeth-Seq (Birkedal et al., 2015; Marchand et al., 2016), which relies on the property of 2’-*O*-methylated riboses to resist alkaline hydrolysis; thus, these modified residues exhibit high ‘cleavage protection’ scores compared to non-methylated sites. Second, a variation of 2OMe-Seq (Incarnato et al., 2017), which exploits the reverse transcription (RT) stops elicited by the 2’-*O*-methylation when using limiting concentrations of deoxyribonucleotide triphosphates (dNTPs); in this method, a higher ratio of RT-stop in low-versus high-dNTP conditions correlates with a higher level of methylation. By combining these two methods, the presence of the three known 2’-*O*-methylated sites in the 16S mt-rRNA was confirmed (**Figure 1A**). Our data also show that MRM1 is responsible for catalysing the ribose methylation of G1145, which is the only position affected by its ablation. Similarly, modification of G1370 is only affected in the absence of MRM3. However, knocking-out either MRM2 or MRM3 impacted on the presence of Um1369, suggesting that this methylation depends on prior modification of G1370 by MRM3. No additional MRM1-, MRM2- or MRM3-dependent 2’-*O*-methylation sites were detected across the mitochondrial transcriptome using these methods (**Figure 1B**).

**Figure 1.**
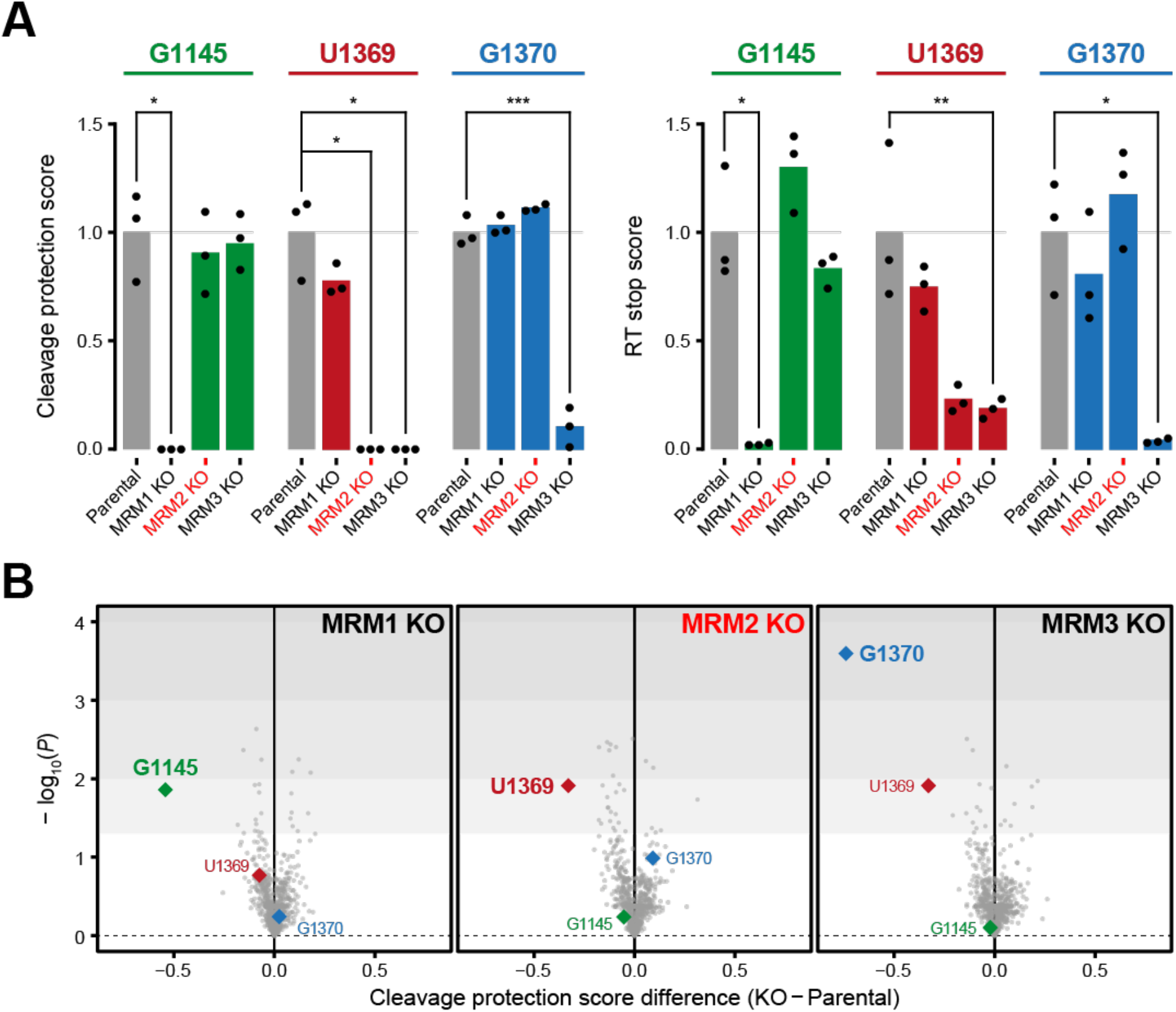
The human mitochondrial transcriptome contains three 2’-*O*-methylated residues, modified by MRM1, MRM2 and MRM3. (A) Determination of ribose methylation levels of known targets of MRM1 (G1145), MRM2 (U1369) and MRM3 (G1370). Experimental values and mean are shown. Statistical significance was assessed using Student’s t-test (∗: *P* ≤ 0.05; ∗∗: *P* ≤ 0.01; ∗∗∗: *P* ≤ 0.001). (B) Mitochondrial transcriptome-wide evaluation of 2’-*O*-methylation in cells devoid of MRM1, MRM2 or MRM3 (*n* = 3 for each cell line). *P* values were determined by Student’s t-test.

### Ablation of MRM2 impairs mitochondrial respiration

Given the association of MRM2 in human mitochondrial pathology (Garone et al., 2017), we focused on understanding its role in mitochondrial function. We first evaluated the functional status of mitochondria in *MRM2* knock-out cells by supplying cells with glucose or galactose as their main carbon source (Gattermann et al., 2004). When cultured in high glucose medium, these cells proliferated at a slower rate, and failed to survive in galactose medium, consistent with a profound and generalised mitochondrial dysfunction (**Figure 2A**). This was corroborated by constitutively high extracellular acidification rate (ECAR, indicative of anaerobic respiration), which was insensitive to inhibition of mitochondrial respiration or uncoupling, and nominal oxygen consumption rate (OCR) (**Figure 2B**). The detected impairment of cellular respiration was characterised in more detail by assessing the activity of mitochondrial respiratory chain (MRC) complexes (**Figure 2C**). While the activity of complex II, the subunits of which are encoded exclusively in the nuclear genome, was not decreased upon depletion of MRM2, that of complexes I and III, which contain mtDNA-encoded core subunits, were significantly decreased (<30% relative to control). Enzymatic activity of complex IV was below detection level and thus it was not possible to accurately determine its value spectrophotometrically. Separation of cellular components in native conditions coupled to in-gel detection of cytochrome c oxidase activity showed that this enzymatic deficiency is due to the virtual absence of assembled complex IV (**Figure S2C**). In agreement with quantitative analysis of the mitochondrial proteome further revealed a reduction in the steady-state levels of MRC components (**Figure 3A**), with the protein presenting the most severe observable down-regulation being MT-CO2, one of the mtDNA-encoded core subunits of complex IV. Taken together, these data show that MRM2 is indispensable for mitochondrial respiratory function in human cells.

**Figure 2.**
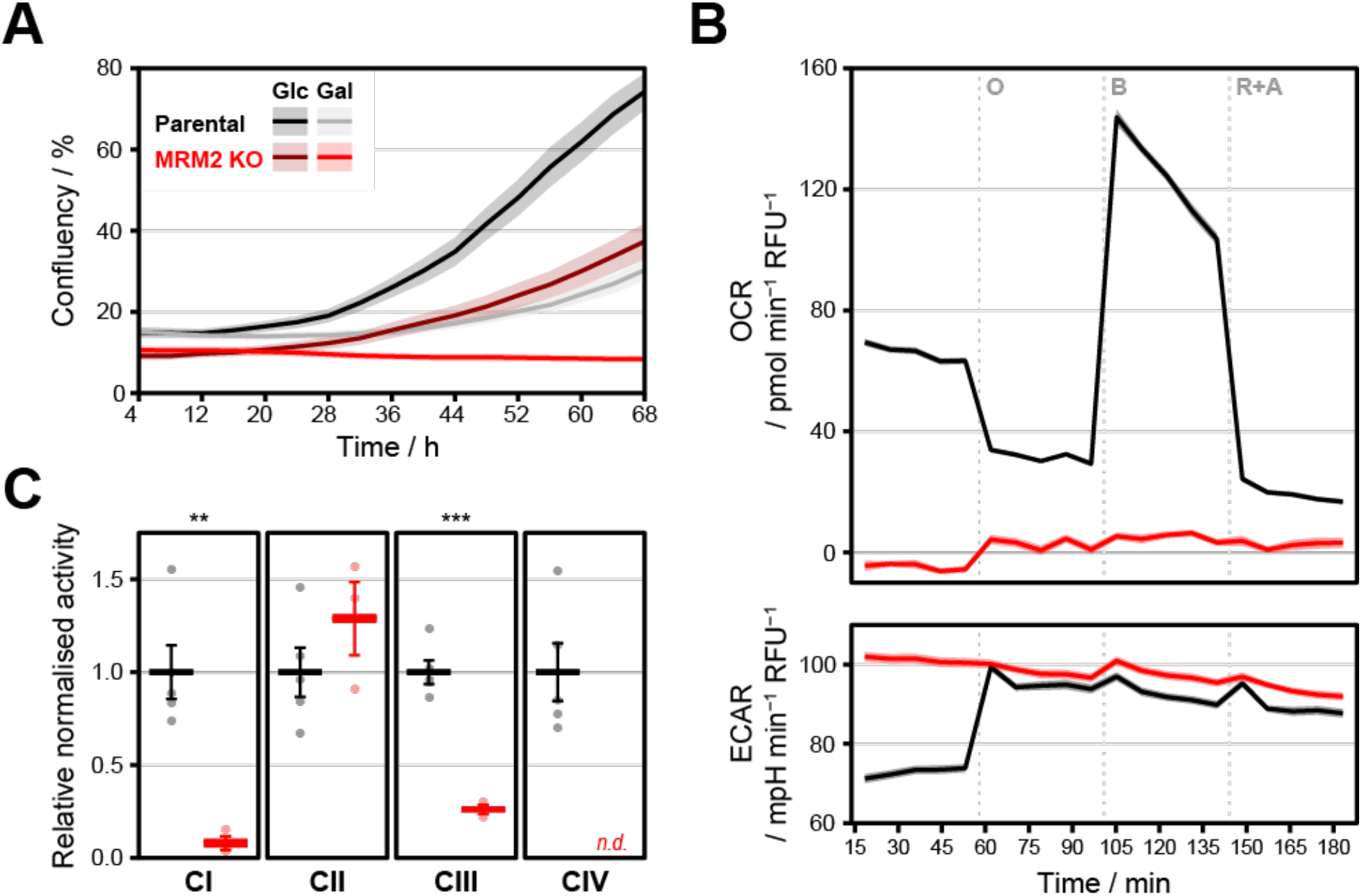
Ablation of MRM2 leads to a severe impairment of cellular respiration and dysfunction of MRC complexes. (A) Cell proliferation in glucose (Glc) or galactose (Gal) media. Data is presented as mean ± SD (*n* = 8). (B) Oxygen consumption rate (OCR) and extracellular acidification rate (ECAR) measurements in the presence of mitochondrial respiration inhibitors (O: oligomycin; B: BAM15; R+A: rotenone and antimycin A). Data is presented as mean ± SD (*n* = 22 for parental, *n* = 23 for *MRM2* knock-out). (C) Spectrophotometric determination of the activity of MRC complexes (CI: complex I, NADH:ubiquinone oxidoreductase; CII: complex II, succinate:ubiquinone oxidoreductase; CIII: complex III, ubiquinol:cytochrome *c* oxidoreductase; CIV: complex IV, cytochrome *c* oxidase). *n.d.*: no data. Experimental values and mean ± SD are shown. Statistical significance was assessed using Student’s t-test (∗∗: *P* ≤ 0.01; ∗∗∗: *P* ≤ 0.001). Data from parental and *MRM2* knock-out (KO) cell lines are represented in black and red, respectively.

**Figure 3.**
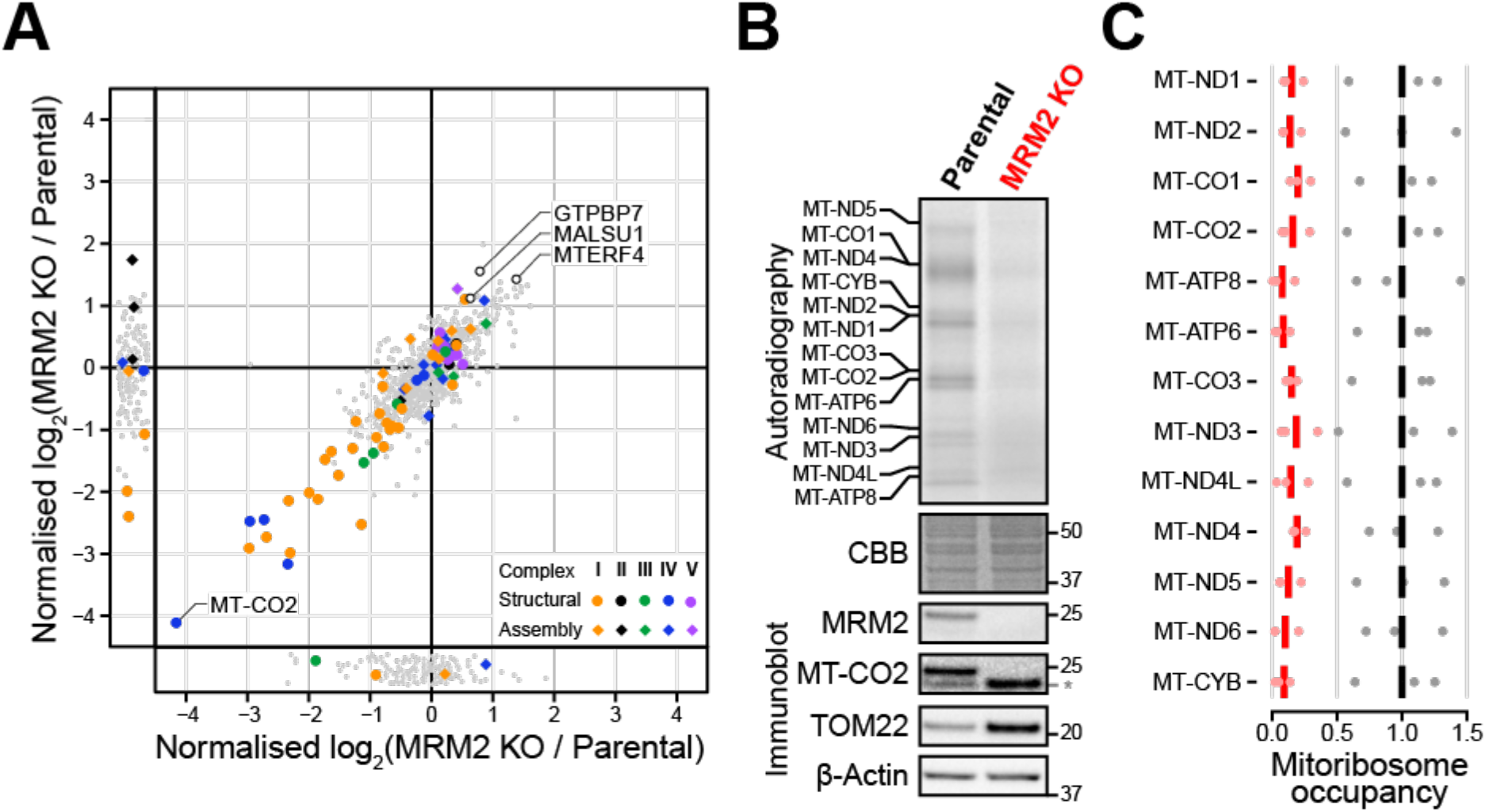
OxPhos complexes are structurally compromised in the absence of MRM2 due to impaired mitochondrial translation. (A) Quantitative mass spectrometric analysis of the mitochondrial proteome of *MRM2* knock-out and parental control cells. Datapoints corresponding to structural components (circles) or assembly factors (diamonds) of OxPhos complexes, and other proteins of interest are highlighted. (B) Metabolic labelling of *de novo* translated mitochondrial proteins (top, quantification in **Figure S2A** and **Figure S2B**) and immunoblotting assessment of steady-state levels of mitochondrial proteins (bottom). Molecular weights of protein standards are presented in kDa to the right of each blot; an asterisk marks a band generated in a previous blot for TOM22. Coomassie brilliant blue (CBB) staining is shown as a loading indicator. (C) Occupancy of mitochondrial transcripts by mitochondrial ribosomes. Experimental values and mean are shown. Data from parental and *MRM2* knock-out (KO) cell lines are represented in black and red, respectively.

### MRM2 is essential for mitochondrial translation

The involvement of MRM2 in 16S mt-rRNA U1369 2’-*O*-methylation, led us to hypothesise that the observed generalised oxidative phosphorylation (OxPhos) defects stem from the perturbation of mitochondrial ribosome-related mechanisms. To test this, the synthesis of mtDNA-encoded proteins was assessed by metabolic labelling of nascent peptides in *MRM2* knock-out and parental cells (**Figure 3B** and **Figure S2A**). The near absence of *de novo* synthesised mitochondrial proteins observed upon MRM2 depletion revealed a considerable dysfunction of mitochondrial translation. The degree to which each mtDNA-encoded protein was affected did not depend on its length (**Figure S2B**), suggesting that this perturbation is not related to translation elongation (Pearce et al., 2017). In order to determine which translation step is affected by the ablation of MRM2, the position of mitochondrial ribosomes along transcripts was determined using high-throughput mitochondrial ribosome footprinting – mitoRibo-Seq (Pearce et al., 2017). Consistent with the previous findings, the quantity of ribosomes engaged in translation was considerably reduced in the absence of MRM2, where the average mitoribosome occupancy on mt-mRNAs was 13.9% ± 7.6% (minimum: 1.6%, median: 12.0%, maximum: 35.5%) of that observed in parental samples (**Figure 3C** and **Figure S3**). No bias in the occupancy of mitoribosomes towards the 5’ or 3’ regions of the coding sequences was observed. Moreover, the pattern of ribosome occupancy along the body of mt-mRNAs was similar between *MRM2* knock-out and parental samples, with no codon-specific stalling. These two pieces of evidence show that a profound impairment of mitochondrial translation underlies the mitochondrial dysfunction phenotype, possibly by perturbing the biogenesis of the mitochondrial ribosome.

### MRM2 is involved in the late stages of mtLSU assembly

To investigate the mechanistic basis for the defect in mitochondrial translation, the composition of mitochondrial ribosomes was assessed using quantitative density gradient analysis by mass spectrometry – qDGMS (Páleníková et al., 2021). While no shifts in the sedimentation profile of mitoribosomal components were observed, proteins of the mtLSU were enriched upon MRM2 depletion (**Figure 4** and **Figure S4**), indicating the accumulation of mature or near-mature forms of this subunit. Given the functional impairment of apparently structurally sound mtLSU particles, these were isolated from *MRM2* knock-out cells and their structure determined to 2.58 Å (**Figure S5**) using cryo-electron microscopy (cryoEM) single particle analysis. Owing to preferential orientation of particles in the specimen, non-tilted and tilted datasets were acquired (**Table S1**). Compared to the mature mtLSU (Brown et al., 2014; Amunts et al., 2015), a large portion of density corresponding to interfacial rRNA is not observed in the consensus map owing to conformational heterogeneity of these elements (**Figure 5A**). This region comprises helices H67-71 (domain IV) and H89-93 (domain V) of the 16S mt-rRNA, encompassing the peptidyl transferase centre (PTC). Additionally, this mtLSU intermediate contains density proximal to uL14m and bL19m corresponding to the MALSU1:L0R8F8:mtACP anti-association module (**Figure 5A** and **Figure S5**). Presence of this module is indicative of particles that are not engaged in translation, as either biogenesis (Brown et al., 2017) or recycling (Desai et al., 2020) intermediates. Since the levels of the recycling factor MTRES1 (C6orf203) in fractions enriched in mtLSU particles from MRM2-depleted mitochondria were comparable to those observed in control cells (**Figure S6**), we interpreted the accumulated mtLSU species as biogenesis intermediates rather than products of defective, damaged or recycling ribosomes.

**Figure 4.**
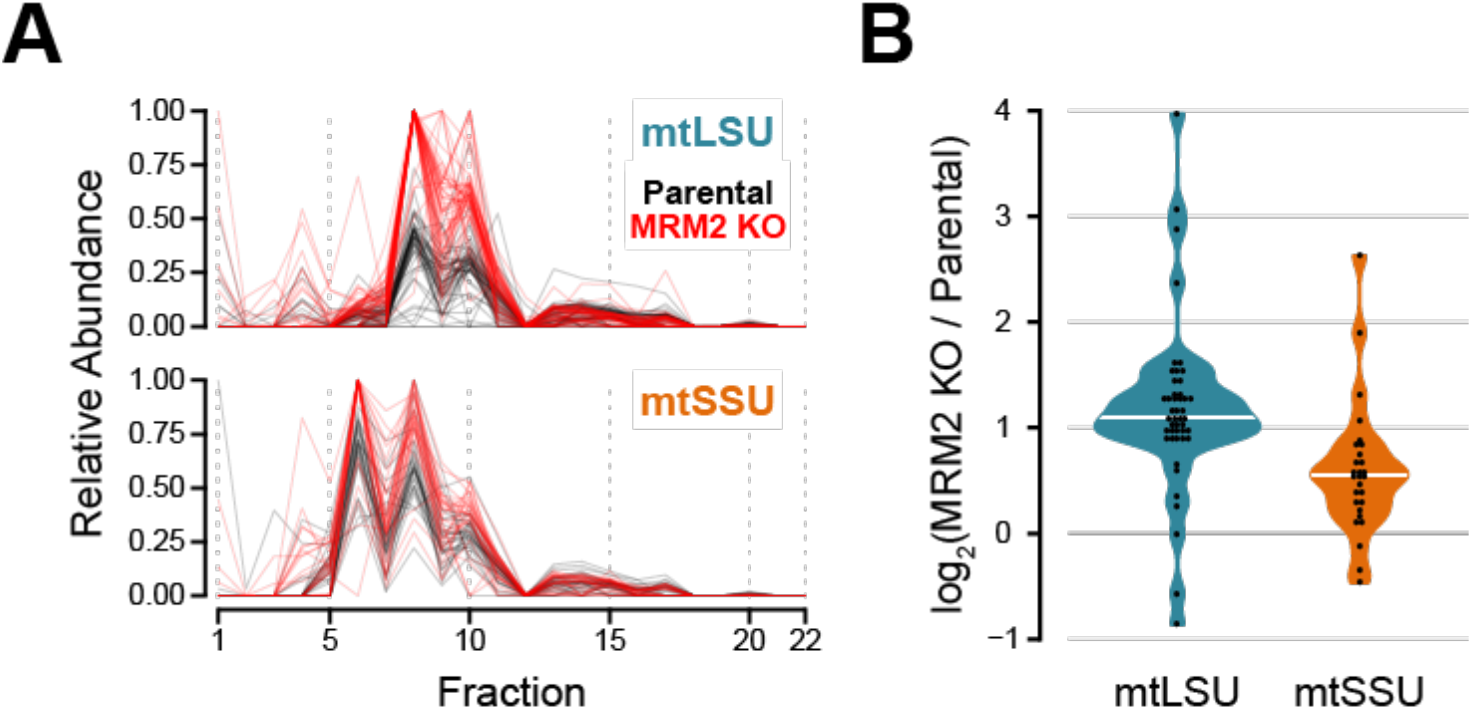
– mtLSU accumulates as mature-like particles in the absence of MRM2. (A) Quantitative gradient fractionation analysis by mass spectrometry (qDGMS) profile of mitoribosomal proteins. A heatmap of individual profiles is shown in **Figure S4**. (B) Assessment of the enrichment of mitoribosomal proteins by subunit. The average from two reciprocal labelling experiments is presented. White horizontal bars represent the median of each distribution.

**Figure 5.**
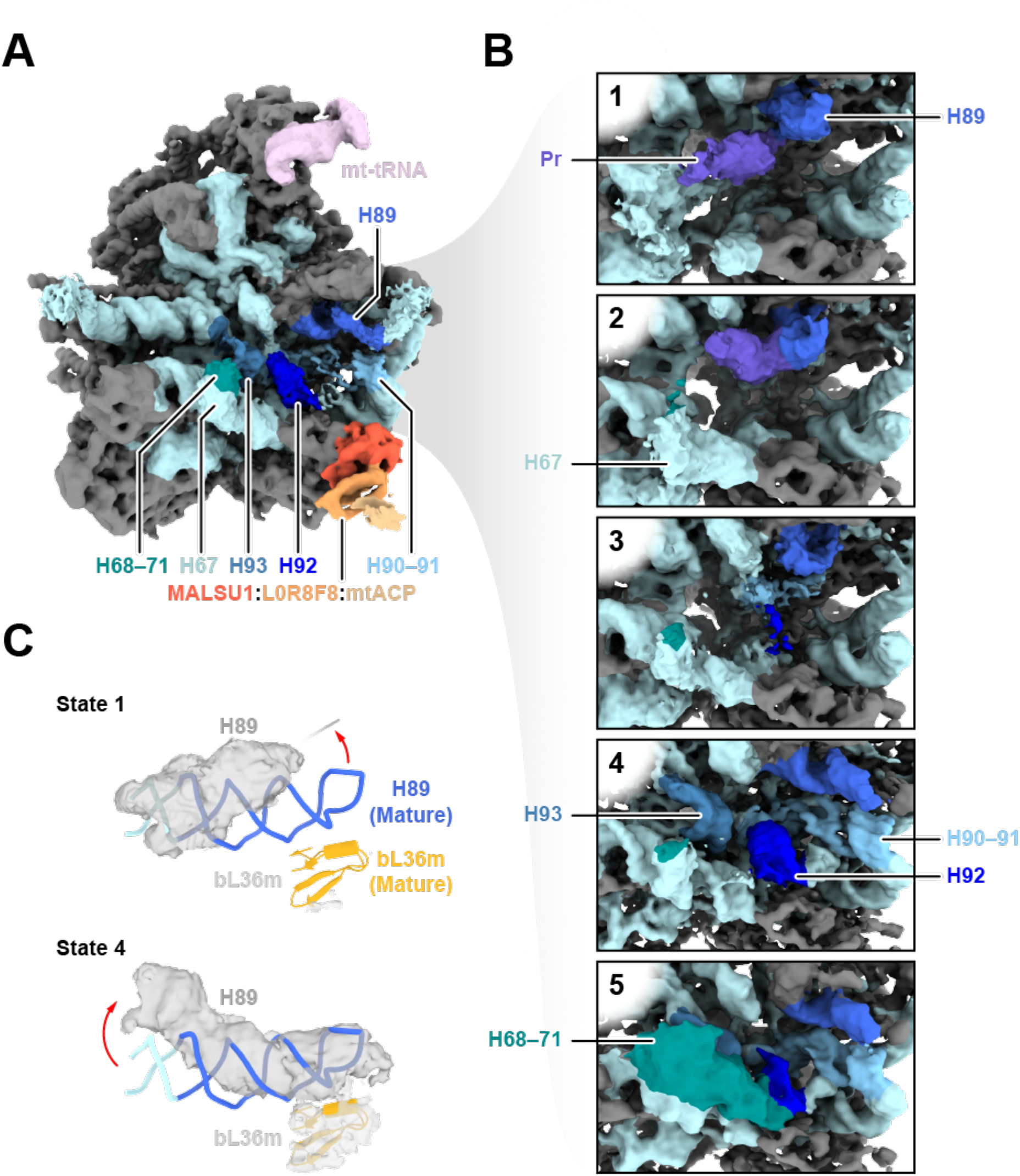
Structure of mtLSU assembly intermediates isolated in the absence of MRM2. (A) Consensus map of the mtLSU intermediate viewed from the intersubunit interface. Relevant features are coloured. MRPs are represented in grey and 16S mt-rRNA in light blue. (B) Clarification of structural heterogeneity of the intersubunit interface by focused classification after masked signal subtraction. States/classes are numbered 1 to 5, corresponding to steps along a putative assembly pathway. Pr: protrusion. Models of relevant regions of states 1 and 4 are presented in **Figure S8A**. (C) Comparison between the conformation of H89 in state 1 (surface), state 4 (surface), and the mature mtLSU (ribbon). Rotation of the apex (state 1) and shift in the base (state 4) of H89 are represented by red arrows. Density corresponding to bL36m in each state is shown as surface. The conformation of H89 and density of bL36m in states 2 and 3 are similar to that modelled in state 1, and those in state 5 are represented by state 4. Density of bL36m is presented for states 1-5 in **Figure S8B**.

Focused classification of the intersubunit interface of these particles showed that this region is highly heterogeneous, with at least five distinct configurations present in the ensemble (**Figure 5B** and **Figure S7**). The most incomplete interface is present in state 1, where H67-71 (domain IV), the apex of H89 (domain V), and H90-93 (domain IV) are not observed. The base of H67 appears in state 2 as a highly disordered region at the base of the intersubunit interface. Furthermore, these two states present density protruding from the PTC, near the base of H89. It is only in state 3 that H92 (where U1369 is located) is slightly stabilised in a near-mature conformation by interaction with H90. In state 4, H89 and H93 become structured and H92 is further stabilised. With the organisation of H89-93, the configuration progresses to state 5, which presents H68-71 in their mature conformation and is thus the most complete and mature-like state; however, this is also the least populated state (∼1% of total number of particles, **Figure S7**), possibly representing particles that were able to stochastically advance through the pathway without the aid of MRM2. Throughout these states, H89 changes from an outwardly rotated conformation with apical flexibility (states 1-3) to one where the whole helix is stabilised along the L7/L12 stalk (states 4 and 5) but its base is shifted to a different position relative to that in the mature state (**Figure 5C**). Concomitantly to H89, the proximal H90-91 are also stabilised in their mature conformation (**Figure S8A**), and bL36m (absent in states 1-3) is present (**Figure S8B**). The MALSU1:L0R8F8:mtACP anti-association module is clearly present in all states and further classification schemes did not produce evidence in favour of the existence of particles lacking this component (**Figure S9A**). To further evaluate structural heterogeneity in the dataset, we used a neural network-based approach – cryoDRGN (Zhong et al., 2021). Using information from whole particles, we corroborated that the intersubunit interface is the main source of structural variability in the studied ensemble of mtLSU particles (**Movie S1**). In addition, this approach revealed particles with density connecting the base of H67 and the PTC, often with an extension to the space between the L1 stalk and the central protuberance (**Figure S8C**). In some cases, including states I and II (states derived from cryoDRGN outputs are indicated in roman numerals), this density could be traced into the back of the PTC, occupying a similar position to the protrusion observed in states 1 and 2. Nonetheless, some density corresponding to H92 is present in state II, providing some evidence that the protrusion does not contain this helix but may be composed of H90, H91, H93 or some combination of these, with an alternative conformation and possibly secondary structure. While H89 is flexible and outwardly rotated in states I and II, it appears in its mature conformation in state III, in which most of the interfacial RNA components (H80-H93) are in mature-like conformations. However, while H68-71 in state III are more ordered than in state 4, they do not reach the same extension as in state 5. Comparing these results with those obtained by focused classification, it was possible to place state I prior to state 3, state II between states 3 and 4, and state III between states 4 and 5.

Additional structural variation was observed in the central protuberance of mtLSU particles from cells devoid of MRM2 (**Figure S9B**). Most particles (∼85%) lack mL40, mL46, mL48 and present a poorly defined structural mt-tRNA, similar to the consensus map. In fewer particles, the structural mt-tRNA appears stabilised and better resolved. Even though poorly defined, density for the missing proteins is observed in ∼7% of the particles. However, this compositional heterogeneity does not reflect in structural variations in other regions of the complex.

Taken together, these data show that MRM2 is involved in the late-stage assembly of mtLSU. However, whether the catalytic activity of MRM2 is required for this role has not been established.

### The catalytic activity of MRM2 is dispensable for mtLSU biogenesis

While acting as an assembly factor of the mtLSU, MRM2 is also a *S*-adenosyl methionine (SAM)-dependent 2’-*O*-ribose methyltransferase with its target on the 16S mt-rRNA. We set out to investigate whether the relevance of MRM2 for mitoribosome biogenesis is due to the chemical modification it introduces in U1369 or to MRM2 acting as a platform for local conformational/compositional changes once bound to assembling mtLSU particles. The putative RNA binding site of MRM2 is composed by evolutionarily conserved surface residues such as K59, which are proximal to the SAM binding site and form an electropositive patch that stabilises the negative charge of the target. The SAM binding pocket is also lined by conserved residues, such as D154, which stabilises the methionine moiety while being positioned close to its Sδ atom (**Figure 6A**; Bügl et al., 2000). Mutation of the catalytic residues K38 and D124 of *Escherichia coli* RlmE (equivalent to *Homo sapiens* MRM2 K59 and D154, respectively) has been shown to abolish its methyltransferase activity (Hager et al., 2002). Validation of the catalytic inactivity of the human variants was obtained by RNA LC-MS^2^, which identified 16S mt-rRNA oligonucleotides containing U1369 solely in its non-methylated state both in *MRM2* knock-out cells complemented with MRM2^K59A^ and MRM2^D154A^, as well as in the knock-out background (**Figure 6B** and **Figure S10**). In control cells, ∼70% of mtLSU-associated 16S mt-rRNA was present as Um1369/Gm1370, with the remainder ∼30% representing the U1369/Gm1370 state; a state lacking the modification deposited by MRM3 was not detected.

**Figure 6.**
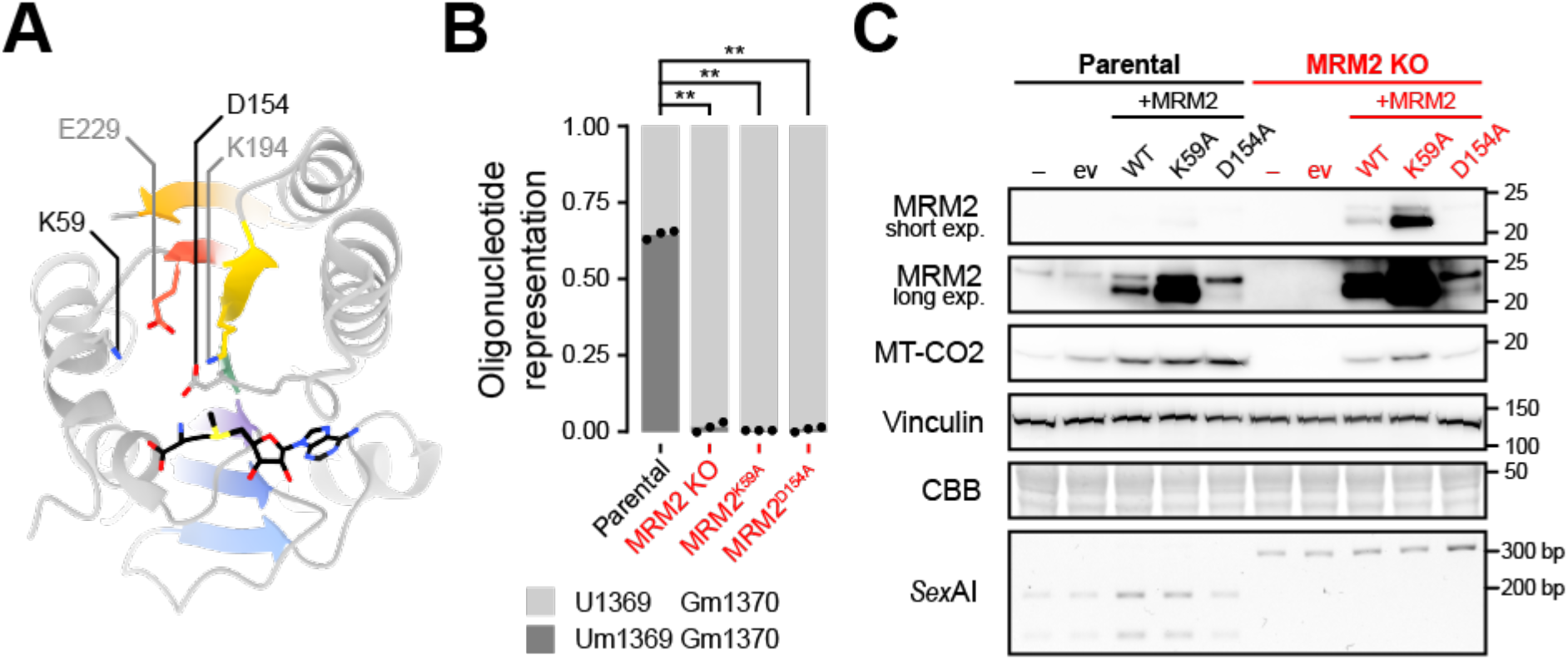
Catalytic mutants of MRM2 are able to restore mitochondrial translation. (A) Structure of human MRM2 (PDB 2NYU; Wu et al., 2006). Evolutionarily conserved active site residues relevant for MRM2-mediated 2’-*O*-methylation are labelled and shown as sticks. *S*-adenosyl methionine (SAM) is represented as black sticks. The seven β-strand methyltransferase fold is coloured by strand, from N- (violet) to C-terminus (red). (B) Quantification of 2’-*O*-methylation of U1369 and G1370 by RNA LC-MS^2^. MRM2^K59A^ and MRM2^D154A^ correspond to samples from MRM2 knock-out cells expressing those catalytic mutants. Statistical significance was assessed using Student’s t-test (∗∗: *P* ≤ 0.01). (C) Immunoblot evaluation of functional rescue of mitochondrial translation in cell lines complemented with wild-type (WT) MRM2, as well as catalytic mutant variants (K59A, D154A). ev: empty vector control. exp.: exposure. Molecular weights of protein standards are presented in kDa to the right of each blot. Coomassie brilliant blue (CBB) staining is shown as a loading indicator. For each cell line, electrophoretically separated *Sex*AI-digested amplicons of the genomic *MRM2* locus targeted for gene editing are presented.

Next, the steady-state level of MT-CO2 was used as a proxy (**Figure 3A** and **Figure 3B**) to assess the functional status of mitochondrial translation (**Figure 6C**). As observed by the restoration of the expression of MT-CO2, both K59A and D154A MRM2 variants were able to rescue the defect in mitochondrial translation caused by knock-out of the endogenous gene despite their null methyltransferase activity. This led to the conclusion that the main contribution of MRM2 to mitochondrial translation is not its methyltransferase activity but its presence and role as an assembly factor, remodelling the intersubunit interface of the assembling mtLSU and thus contributing towards the mature conformation.

### Other mtLSU biogenesis factors are not redundant to MRM2

Overexpression of certain ribosomal biogenesis factors has been reported to rescue the severe growth and 50S biogenesis defects of bacteria lacking the orthologue of MRM2 (RlmE/RrmJ/FtsJ/MrsF). This was the case of the GTPases ObgE/CgtA and EngA (Tan et al., 2002). GTPases have been implicated in the biogenesis of the human mitochondrial ribosome, some of which orthologous to bacterial proteins. To test whether mitochondrial GTPases can rescue the MRM2 ablation phenotype in human, we complemented *MRM2* knock-out cells with GTPBP5/MTG2, GTPBP7/MTG1, and GTPBP10, which are known to be present in mitochondria and participate in mtLSU biogenesis (Lavdovskaia et al., 2018; Maiti et al., 2018; Lavdovskaia et al., 2020; Maiti et al., 2020; Cipullo et al., 2021). However, despite the overexpression of these proteins, mitochondrial translation was not functionally restored, as indicated by the absence of MT-CO2 in the complemented cells (**Figure S11**), indicating a nonredundant role of MRM2 in the assembly of mtLSU.

### MRM2 is required for organismal homeostasis

Given the severe phenotype caused by ablation of MRM2 in cultured cells, as well as reports of patients harbouring variants in the coding gene, presenting with a Mitochondrial Encephalomyopathy, Lactic Acidosis, and Stroke-like episodes (MELAS)-like pathology with progressive encephalopathy and stroke-like episodes (Garone et al., 2017), we investigated the relevance of this protein for whole-organism and tissue-specific homeostasis in a *Drosophila melanogaster* model. Therefore, we knocked-down the *MRM2 D. melanogaster* orthologue *CG11447* (henceforth referred to as *DmMRM2*) by expressing an inducible (UAS-) RNAi construct driven by the ubiquitous promoter *da-GAL4* driver (**Figure 7A**). Downregulation of DmMRM2 led to developmental delay, with few larvae progressing to pupae, with most of these dying at late pupal stage. In rare cases where eclosion was successful, adults were either trapped by the pupal case while emerging or immobilised on the food as they were too weak to escape. These adults showed anterior thoracic indentations alongside deformed wings and flattened abdomen (**Figure 7B**). *DmMRM2* knock-down was lethal when performed under another ubiquitous driver (*act5C-GAL4*) or a pan-neuronal driver (*nSyb-GAL4*), further indicating the essentiality of this protein.

**Figure 7.**
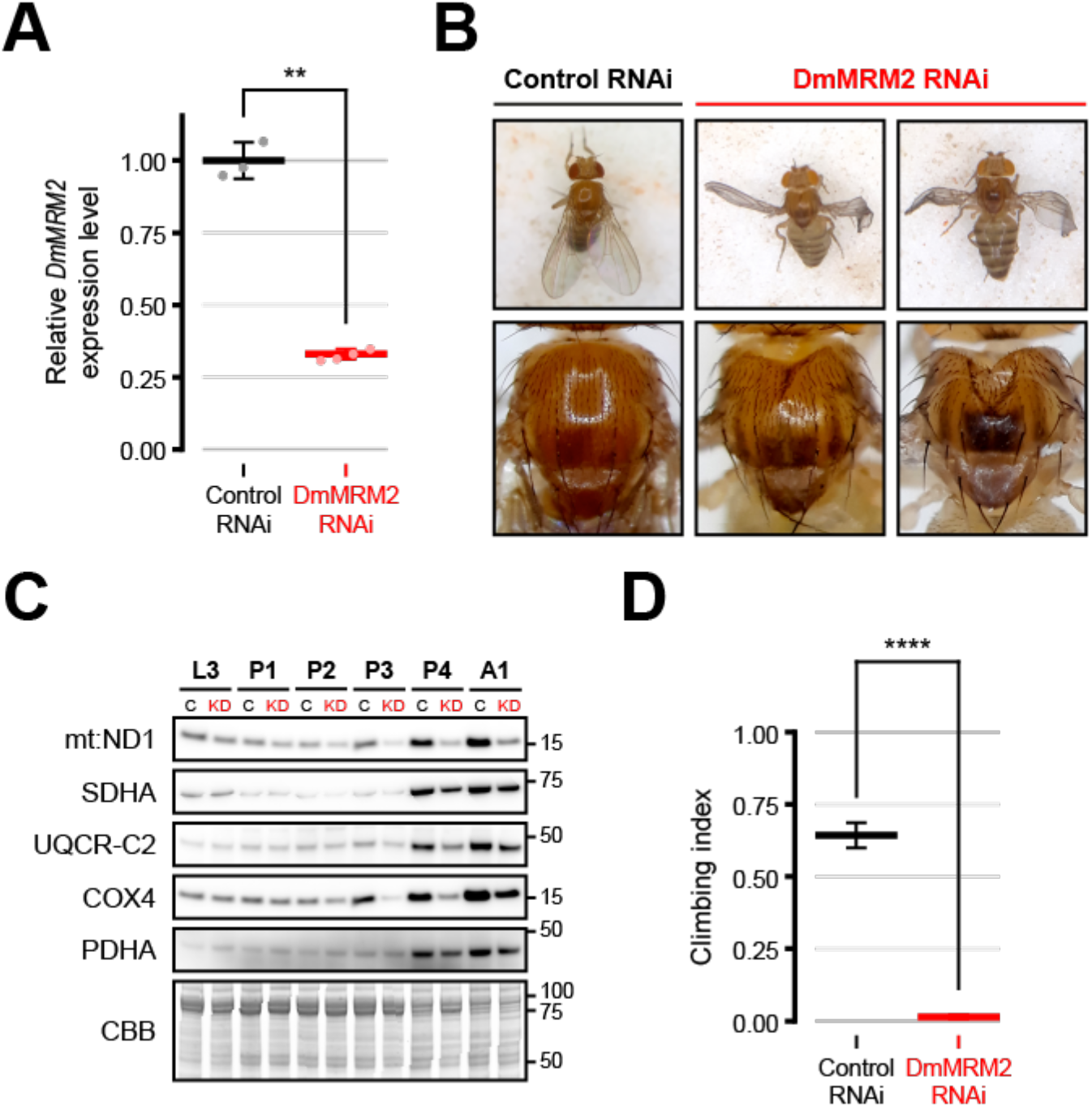
DmMRM2 is detrimental for the development of *D. melanogaster*, in particular towards the end stages of pupation. (A) Levels of *DmMRM2* transcripts in L3 larvae ubiquitously expressing control (*n* = 3) or DmMRM2 (*n* = 4) RNAi under the *da-GAL4* driver. Values are normalised by the level of *αTub84B* transcripts. Data is presented as individual datapoints and mean ± SD. Each datapoint was generated from three L3 larvae. Statistical significance was assessed using Student’s t-test (∗∗: *P* ≤ 0.01). (B) Dorsal view of whole-body and thorax of control and DmMRM2 knock-down adult flies. (C) Immunoblot assessment of the steady state levels of MRC subunits over the course of pupation in control (C) and *DmMRM2* knock-down (KD) individuals. L3: third instar larvae; P1: day 1 pupae; P2: day 2 pupae; P3: day 3 pupae; P4: day 4 pupae; A1: adults after eclosion. Molecular weights of protein standards are presented in kDa to the right of each blot. Coomassie brilliant blue (CBB) staining is shown as a loading indicator. (D) Startle-induced negative geotaxis (climbing) assay performed on adult male flies expressing control (*n* = 70) or *DmMRM2* (*n* = 56) RNAi under the pan-muscular *Mef2-GAL4* driver. Data is presented as mean ± SEM. Statistical significance was assessed using the Kruskal-Wallis nonparametric test (∗∗∗∗: *P* ≤ 0.0001). Control RNAi: *UAS-lacZ RNAi; da-GAL4* (transcript quantification, imaging, immunoblotting), *UAS-lacZ RNAi; Mef2-GAL4* (climbing assay).

To evaluate the correlation between the lethal phase and requirements on mitochondrial function, the steady-state level of subunits of MRC complexes was assessed in different stages of development (**Figure 7C**). Third instar larvae (L3) and young adults contrasted in the amount of these subunits, which correlates with their different metabolic needs (Da-Ré et al., 2014; Beebe et al., 2020). This cellular and molecular shift occurred during pupation, where the steady-state levels of MRC subunits were progressively upregulated. Steady-state levels increased considerably during the second half of pupation, suggesting a stronger reliance of cells on mitochondrial respiration. This increase was less marked in DmMRM2 knock-down animals, with levels of MRC subunits in late pupation stages being considerably lower when compared to matched control animals. Concurrently, the development of DmMRM2 knock-down individuals is arrested at this stage, linking the observed lethality to the role of this protein in mitochondrial function and its relevance for cell and organismal homeostasis.

In order to investigate the impact of these molecular alterations at the whole organism level, and since muscular tissue is commonly affected in mitochondrial disorders (Gorman et al., 2016), including the MELAS-like presentation of patients harbouring MRM2 mutations (Garone et al., 2017), *DmMRM2* was knocked-down using a pan-muscular driver (*Mef2*-GAL4). While this did not lead to pupal lethality, adult flies presented considerably impaired locomotor ability in the startle-induced negative geotaxis assay, which monitors neuromuscular function (**Figure 7D**). Taken together these data show that MRM2, through its involvement in mtLSU biogenesis, is indispensable for organismal homeostasis.

## DISCUSSION

To understand the role of proteins associated with human pathogenesis, it is necessary to comprehend their function and the molecular mechanisms in which they are involved. In this study, we performed a multidisciplinary analysis of the mitochondrial role of MRM2, a protein that underlies the molecular basis of the MELAS-like syndrome in patients harbouring variants of the coding gene.

We demonstrate that all known 2’-*O*-methylations in human mitochondrial transcripts, G1145, U1369 and G1370, are introduced by MRM1, MRM2 and MRM3, respectively, and using transcriptome-wide approaches we detect no additional targets for these enzymes in mtRNAs (**Figure 1**). As mentioned, MRM2 has been associated with human disease with a typical mitochondrial cytopathy presentation (Garone et al., 2017); to study this in greater detail, we first generated a cellular knock-out model. Ablation of MRM2 leads to mitochondrial dysfunction with severe reduction in cellular respiration and activity of MRC complexes containing mitochondrially-encoded subunits (**Figure 2**). We show that this is caused by the impairment of mitochondrial translation, leading to reduced production of mtDNA-encoded proteins (**Figure 3**, **Figure S2** and **Figure S3**). Consequently, this impacts on the structural integrity of the OxPhos system and thus explains its functional collapse. The defect in mitochondrial translation is associated with the large subunit of the mitochondrial ribosome, which RNA core element (16S mt-rRNA) is 2’-*O*-methylated by MRM2 in the A-loop residue U1369. Despite shown to be dysfunctional in the absence of MRM2, mtLSU particles contain virtually all of their MRPs and accumulate in a near mature state (**Figure 4** and **Figure S4**).

The late stages of mtLSU assembly involve almost exclusively the remodelling of interfacial RNA elements of otherwise complete particles, as shown by recent structural studies (Chandrasekaran et al., 2021; Cheng et al., 2021; Cipullo et al., 2021; Hillen et al., 2021; Lenarčič et al., 2021). Key players include the helicase DDX28, which dislocates the central protuberance to increase solvent exposure of the intersubunit interface, the MTERF4:NSUN4 complex, which holds H68-71 of domain IV of 16S mt-rRNA in an immature conformation to allow access of assembly factors to the PTC, and the GTPases GTPBP5, GTPBP6, GTPBP7 and GTPBP10, which coordinate the maturation of the PTC through a series of conformational rearrangements. In addition to these, the MALSU1:L0R8F8:mtACP module is bound in the reported assembly intermediates, avoiding their premature association with mtSSU particles and preventing the immature mtLSU from engaging in translation (Brown et al., 2017).

In the absence of MRM2, there is an accumulation of mtLSU particles containing the MALSU1 anti-association module and unstructured interfacial RNA components, similar to homeostatic assembly intermediates reported previously (**Figure 4** and **Figure 5**; Brown et al., 2017). However, in contrast to homeostatic conditions, where these intermediates correspond to a relatively small portion of the observed mtLSU particles, this representation is virtually absolute in cells devoid of MRM2. Based on the features of the obtained intermediates, we hypothesise the following sequence of assembly steps, starting from a state lacking bL36m, with unstructured H67-71 (domain IV) and H90-93 (domain V, possibly in an alternative RNA structure): (1) stabilisation of the base of the intersubunit interface via H67; (2) initial folding of H92 by interaction with the base of H90; (3) folding of H90-93, inward rotation of H89 by interaction with H91 and concomitant structuring of the bL36m binding pocket, favouring its incorporation; (4) refolding of H68-71 via interaction with stabilised interface components, including H92. In the absence of MRM2, most particles accumulate before step (3) can occur, indicating that this protein contributes to the late stages of mtLSU biogenesis not only by stabilising its binding substrate, H92, in the proper folding configuration, but also by relocating H89 and H91. Although maturation of domain IV does not seem strictly dependent on the presence of MRM2, as evidenced by a low number of particles where these components are structured (**Figure 5B** state 5, **Figure S7**), it may be catalysed by the folding of proximal RNA helices and the presence of Um1369, which participates in the bacterial Um2552 (H92) – C2556 (H92) – U1955 (H71) interaction triad (Arai et al., 2015), corresponding to Um1369 (H92) – C1373 (H92) – U948 (H71) in human mtLSU, and may contribute to the positioning of G1370 by steric hindrance created by the 2’-methoxy of Um1369.

Analysis of residual heterogeneity in the dataset (**Figure S8C** and **Movie S1**) brought to light additional states of mtLSU particles, some of which recapitulate results obtained by conventional focused classification with signal subtraction (FCwSS), while others probe additional configurations in the mtLSU conformational space and/or assembly pathway. This enabled a clearer visualisation of density of unknown identity, with RNA helix-like features, protruding from the PTC and in proximity of the unstructured H68-71. Furthermore, it added to the evidence against the existence of an appreciable number of mtLSU particles without the MALSU1:L0R8F8:mtACP module in the absence of MRM2.

It was shown that the bacterial ribosomal protein bL36 integrates into LSU particles late during assembly, which is triggered by the presence of the MRM2 orthologue, RlmE/RrmJ/FtsJ/MrsF (Arai et al., 2015). Inspection of the mtLSU assembly intermediates described in the present work shows that incorporation of bL36m is MRM2-independent (**Figure 5** and **Figure S8**), and most likely relies more directly on the acquisition of a proper folding of interfacial rRNA elements, namely H89, which in turn depends on the folding of H91 via stabilisation of H92, aided by MRM2.

Moreover, the final stages of the assembly of mtLSU particles are independent of the 2’-*O*-methylation of U1369 by MRM2 (**Figure 6**). However, they depend on the transit through conformational states aided by a set of proteins, including MRM2. In the absence of this protein, mtLSU particles are still able to stochastically reach a near mature conformation, even though at a nominal rate. Nevertheless, since the conformation of these mtLSU is not similar enough to that seen in mature particles (displaced H89 base), they are not licenced to engage in translation, as seen by the persistence of the MALSU1:L0R8F8:mtACP module. Thus, the strict quality control of mtLSU assembly is fulfilled. This adds to previous evidence that the binding of assembly factors with RNA modification activity to assembling cytosolic ribosomes is frequently associated to quality-control checkpoints during the biogenesis of these complexes. However, the relevance of those enzymes, often evolutionarily conserved, is not related to the modification they introduce in rRNAs, but to their presence in cells (Sharma and Lafontaine, 2015).

Analysis of the methylation of 16S mt-rRNA nucleotides in *MRM1-3* knock-out models showed that the 2’-*O*-methylation of U1369 by MRM2 depends on the prior modification of U1370, substrate of MRM3 (**Figure 1**). This implies that MRM3 exerts its activity over the assembling mtLSU prior to MRM2, adding molecular evidence to support structural findings (Chandrasekaran et al., 2021; Cheng *et al*., 2021; Cipullo et al., 2021; Hillen et al., 2021; Lenarčič et al., 2021).

In bacterial systems, ablation of the MRM2 orthologue can be compensated by the overexpression of certain GTPases (ObgE/CgtA and EngA). The resulting bacterial ribosomal large subunits still lack 2’-*O*-methylated U2552 (equivalent to U1369 in human mitochondria) but are nonetheless able to engage in translation (Tan *et al*., 2002). In *S. cerevisiae*, only a partial suppression in the thermosensitive loss of mtDNA of a *Δmrm2* mutant strain was observed when a eukaryotic orthologue of ObgE, Mtg2p (MTG2/GTPBP5 in human), was overexpressed (Tan *et al*., 2002; Datta, Fuentes and Maddock, 2005). Although expression of human orthologues of mitochondrial GTPases did not complement the knock-out of *MRM2* (**Figure S11**), catalytic mutants of this methyltransferase were able to functionally restore mitochondrial translation without recovering the modification of U1369 (**Figure 6**), a process for which the mitochondrial system seems to lack redundancy.

We further tested the role of MRM2 in the model organism *D. melanogaster*, showing that its orthologue DmMRM2 is detrimental for development. This is especially relevant upon the shift from a larval glycolytic phenotype that supports rapid growth, to a more efficient respiratory phenotype in adults, which occurs during pupation (Da-Ré et al., 2014; Beebe et al., 2020). It is possible that maternal deposition of factors and mitochondria contribute to the maintenance of mitochondrial function during larval stages when *DmMRM2* is knocked-down. However, the turnover of OxPhos complexes, unmet by the lower rate of synthesis of their core subunits, as well as the increased reliance on OxPhos itself eventually tip the homeostatic balance during pupation towards an unbearable cellular burden (**Figure 7**). Furthermore, *DmMRM2* knock-down in *D. melanogaster* has the potential to be a model of mitochondrial dysfunction that does not target the OxPhos system directly but broadly affects its mtDNA-encoded components. This constitutes a useful tool for further investigation of fundamental questions, such as those related to differential effects of mitochondrial dysfunction associated with tissue specificity.

Our results identify a crucial point of control during the assembly of the mammalian mtLSU with mechanistic insights that enlighten not only the role of MRM2 in mitochondrial translation, but also the phenotype caused by its disruption, and consequent implications at the multi-tissue level. Given the contraction in the number of modified residues in the mammalian mitochondrial ribosome and the knowledge on the enzymatic repertoire responsible for these modifications, it will be important to determine the role and essentiality of the remaining modifications and/or the involved enzymes in order to better understand mitoribosome biogenesis and mitochondrial translation.

---

**Figure S1.**
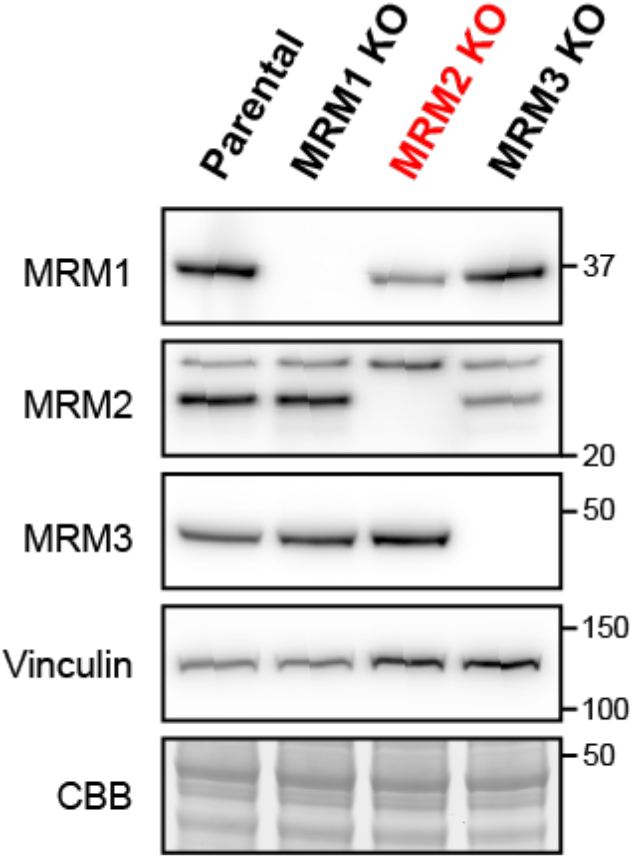
Validation of the MRM1, MRM2, and MRM3 knock-out cell lines. Immunodetection of the three mitochondrial 2’-*O*-methyltransferases in cellular lysates from parental and each of the knock-out cell lines. Molecular weights of protein standards are presented in kDa to the right of each blot. Coomassie brilliant blue (CBB) staining is shown as a loading indicator.

**Figure S2.**
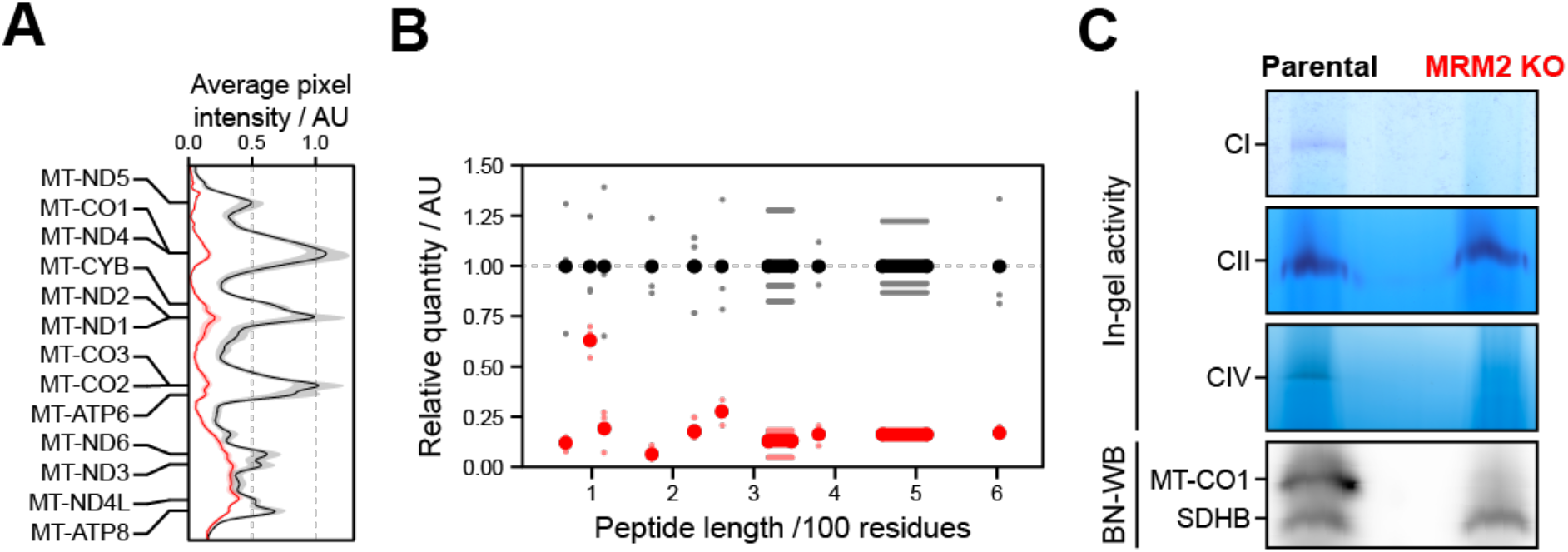
Investigation of synthesis, assembly and activity of OxPhos complexes in the absence of MRM2. (A) Average profile of mtDNA-encoded proteins observed in Figure 3B (*n* = 3). Traces from parental and MRM2-depleted samples are presented in black and red, respectively. (B) Quantification of mtDNA-encoded proteins from metabolic labelling (Figure 3B) plotted against their peptide length. Datapoints (small dots) and corresponding mean values (*n* = 3, larger solid dots) representing samples from parental and MRM2-depleted cells are presented in black and red, respectively. (C) NADH oxidase (CI), succinate oxidase (CII) and cytochrome *c* oxidase (CIV) activities evaluated in cell extracts. Steady-state levels of complexes IV (MT-CO1) and II (SDHB) were evaluated by immunoblotting of cellular contents separated in near-native conditions.

**Figure S3.**
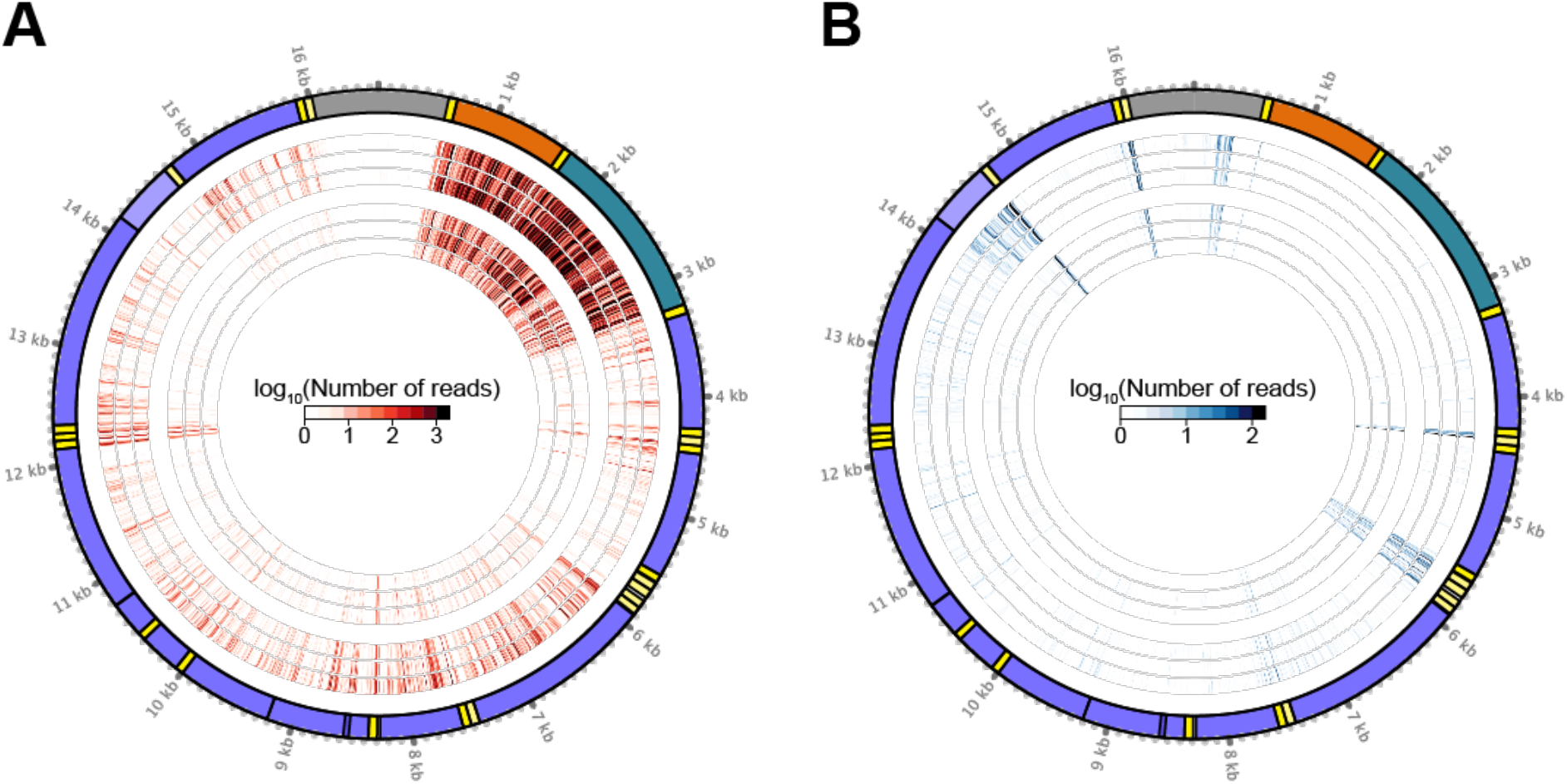
Distribution of ribosomal occupancy along mitochondrial transcripts. Overview of the mitochondrial ribosome footprints mapped to the mitochondrial reference sequence. Heavy-strand promoter- (A) and light-strand promoter- (B) derived reads. The 5’ read ends from the triplicate parental samples are presented in the three outer rings and those from MRM2 knock-out samples are presented in the three inner rings. The mitochondrial transcriptome is represented in a circular plot to facilitate visual comprehension. Yellow: mt-tRNAs, purple: mt-mRNAs, orange: 12S mt-rRNA, teal: 16S mt-rRNA, grey: D-loop. Transcripts encoded in the anti-sense strand are represented in the circular ideogram as lighter crown arcs. Mitoribosome occupancy is presented for each mitochondrial transcript in Figure 3C.

**Figure S4.**
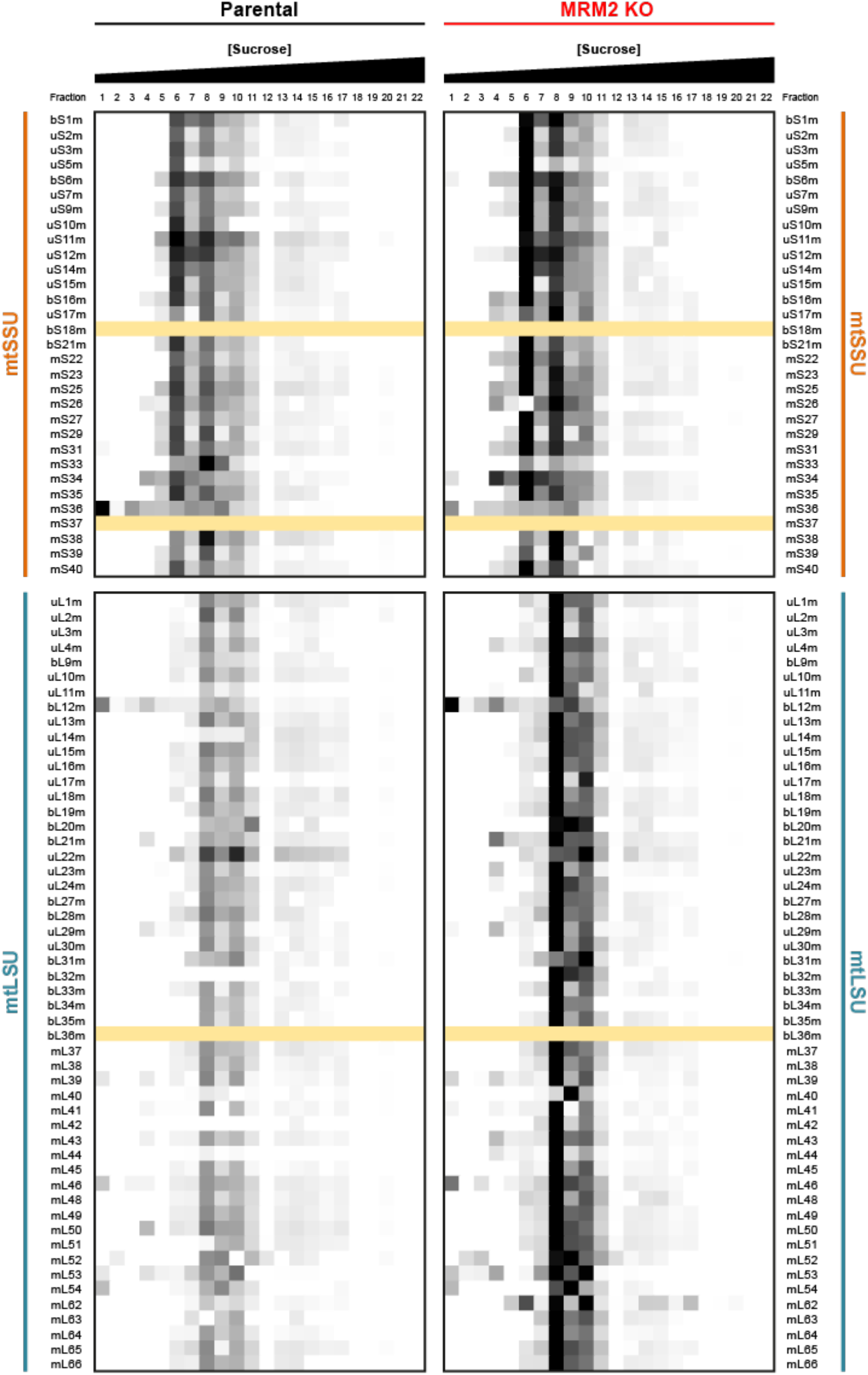
Proteomic characterisation of mitoribosome components. Heatmap of the quantitative profile of mitoribosomal proteins in fractions collected from a continuous sucrose density gradient where mitochondrial extracts were resolved. The profile of each detected protein is coloured according to their abundance in each fraction (low to high abundance is coloured from white to black, respectively). Proteins for which no peptides were detected in parental and *MRM2* knock-out samples are shown in yellow. A summary of these results is presented in Figure 4.

**Figure S5.**
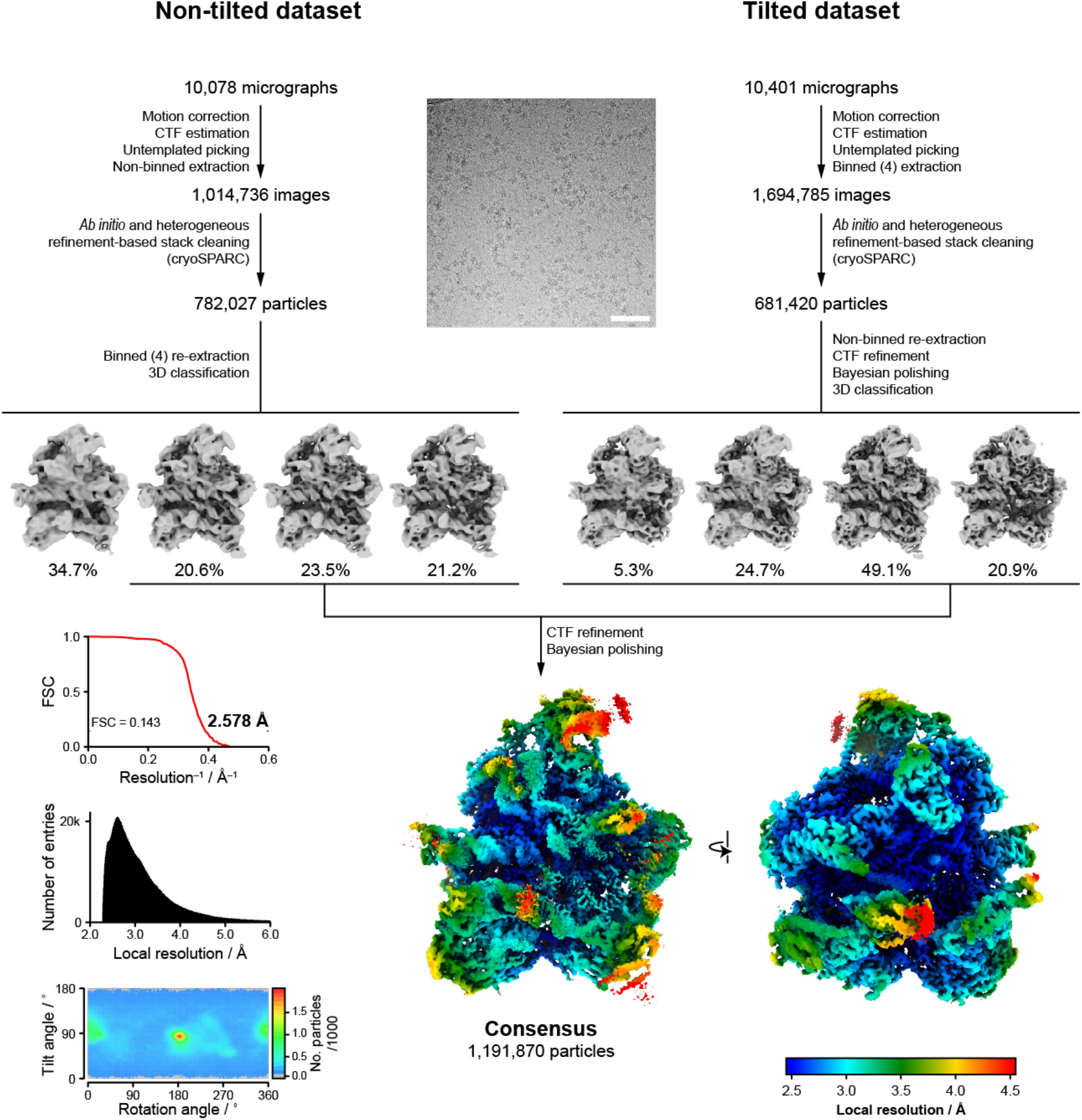
CryoEM data processing workflow. Processing strategy is presented for the collected datasets when worked separately, and after being merged. An example micrograph is shown (scale bar: 100 nm), as well as the outcome maps of key classification steps (particle distribution is presented as percentages below the map of each class) and final consensus maps (coloured by local resolution). Fourier shell correlation (FSC), local resolution distribution and angular distribution are presented in their respective plots. Related to Figure 5A.

**Figure S6.**
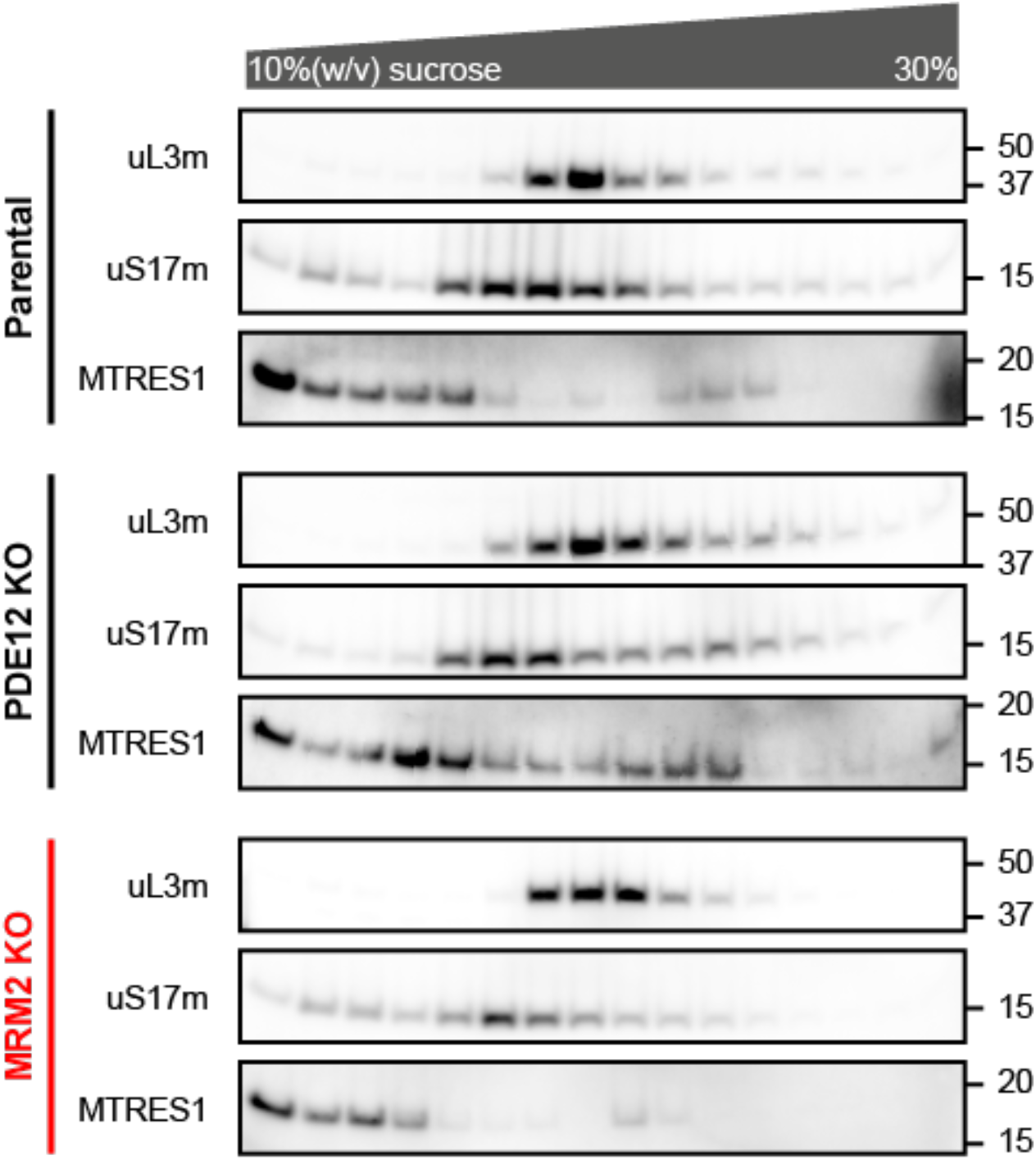
Investigation of the presence of mtLSU-associated recycling factors. Immunoblot profile of mtLSU (uL3m), mtSSU (uS17m) and the recycling factor MTRES1 along a continuous density gradient. Molecular weights of protein standards are presented in kDa to the right of each blot.

**Figure S7.**
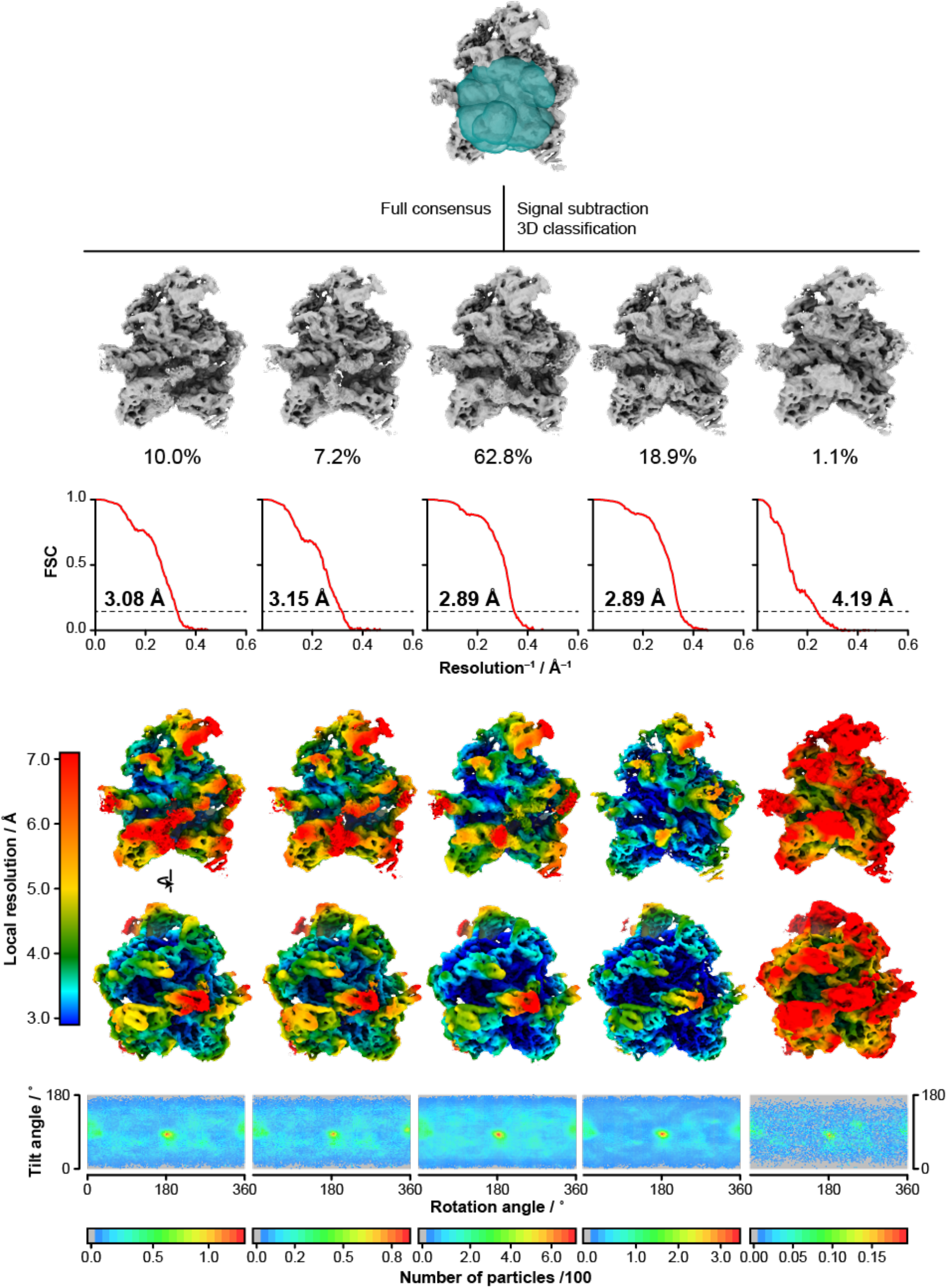
Details of the focused classification on the intersubunit interface. A mask containing the intersubunit interface (teal surface) was used to perform signal subtraction. Classification (T=300) of the resulting signal generates the five presented classes (state 1 to 5 from left to right; particle distribution is presented as percentages below the map of each class). Fourier shell correlation (FSC), surface colouring by local resolution (top: intersubunit interface view; bottom: 180° turn, peptide exit tunnel view), and angular distribution are presented. Overall resolution of each map is presented for FSC=0.143 (dashed line). Related to Figure 5B.

**Figure S8.**
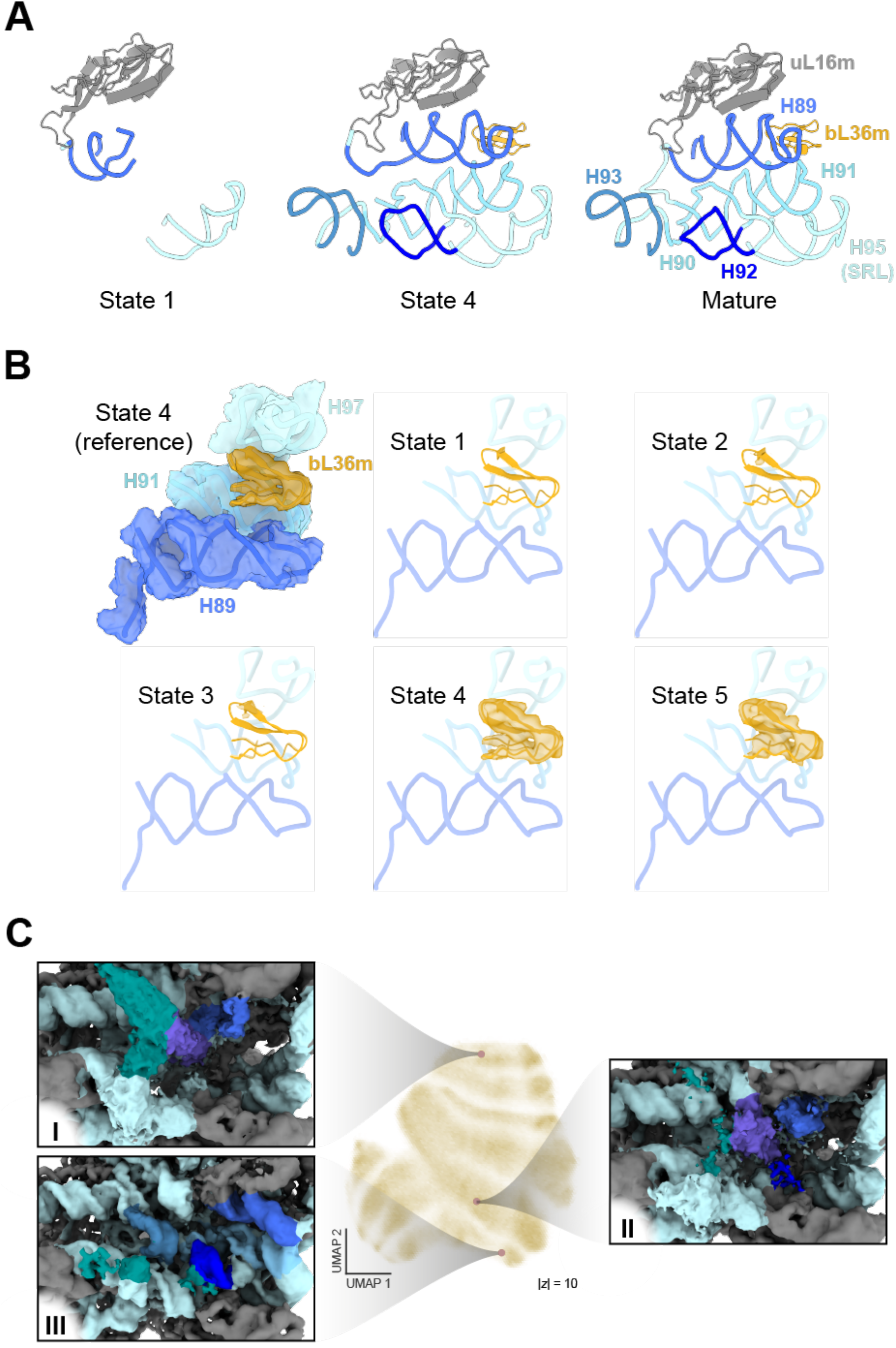
Inspection of heterogeneity of mtLSU intersubunit interface components. (A) Conformational and compositional heterogeneity of H89-93, H95 (containing the sarcin–ricin loop, SRL) and associated ribosomal proteins. Comparison of the modelled assembly intermediate states with the structure of the mature mtLSU (PDB 3J9M). (B) Evaluation of bL36m density (dark orange surface) across the different mtLSU conformation states. The model of state 4 is shown for all states merely as a guide for the expected (states 1-3) or actual (states 4 and 5) location of H89 and bL36m. H91 (domain V) and H97 (domain IV) are shown as proximal interactors of bL36m in the L7/L12 stalk. (C) Recapitulation of structural intermediates using a neural network approach (cryoDRGN). Uniform manifold approximation and projection (UMAP) representation of the latent space is shown (centre), alongside highlighted representations of selected cluster centres (left and right). Latent space exploration is shown in greater extension in **Movie S1**. Surfaces and ribbons are presented with the same colour code as Figure 5A.

**Figure S9.**
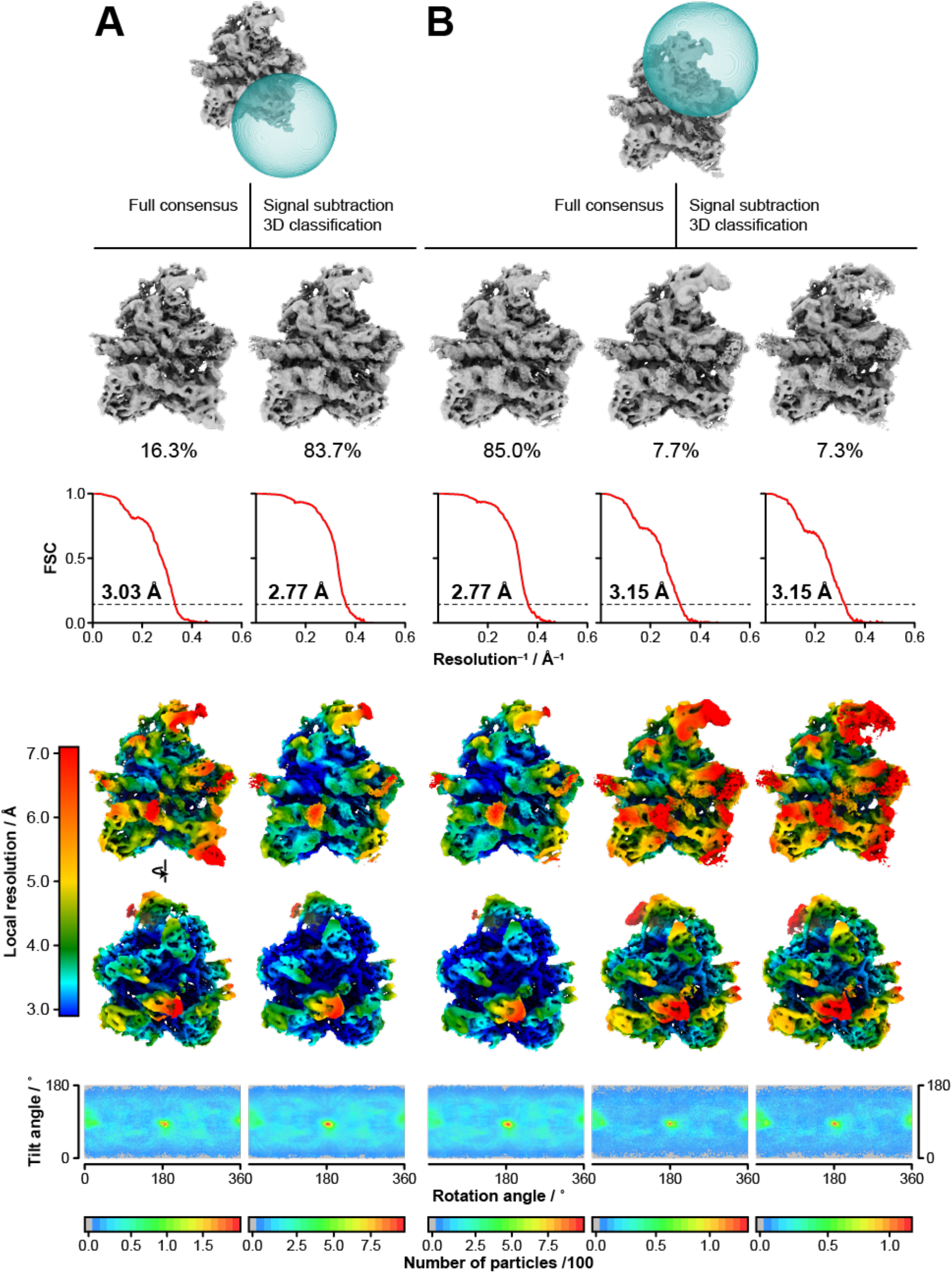
Details of the focused classification on additional mtLSU regions. Masks (teal surface) containing (A) the MALSU1:L0R8F8:mtACP module or (B) the central protuberance were used to perform signal subtraction. Classification (T=300 for MALSU1 module, T=50 for central protuberance) of the resulting signal generates the presented classes (particle distribution is presented as percentages below the map of each class). Fourier shell correlation (FSC), surface colouring by local resolution (top: intersubunit interface view; bottom: 180° turn, peptide exit tunnel view), and angular distribution are presented. Overall resolution of each map is presented for FSC=0.143 (dashed line).

**Figure S10.**
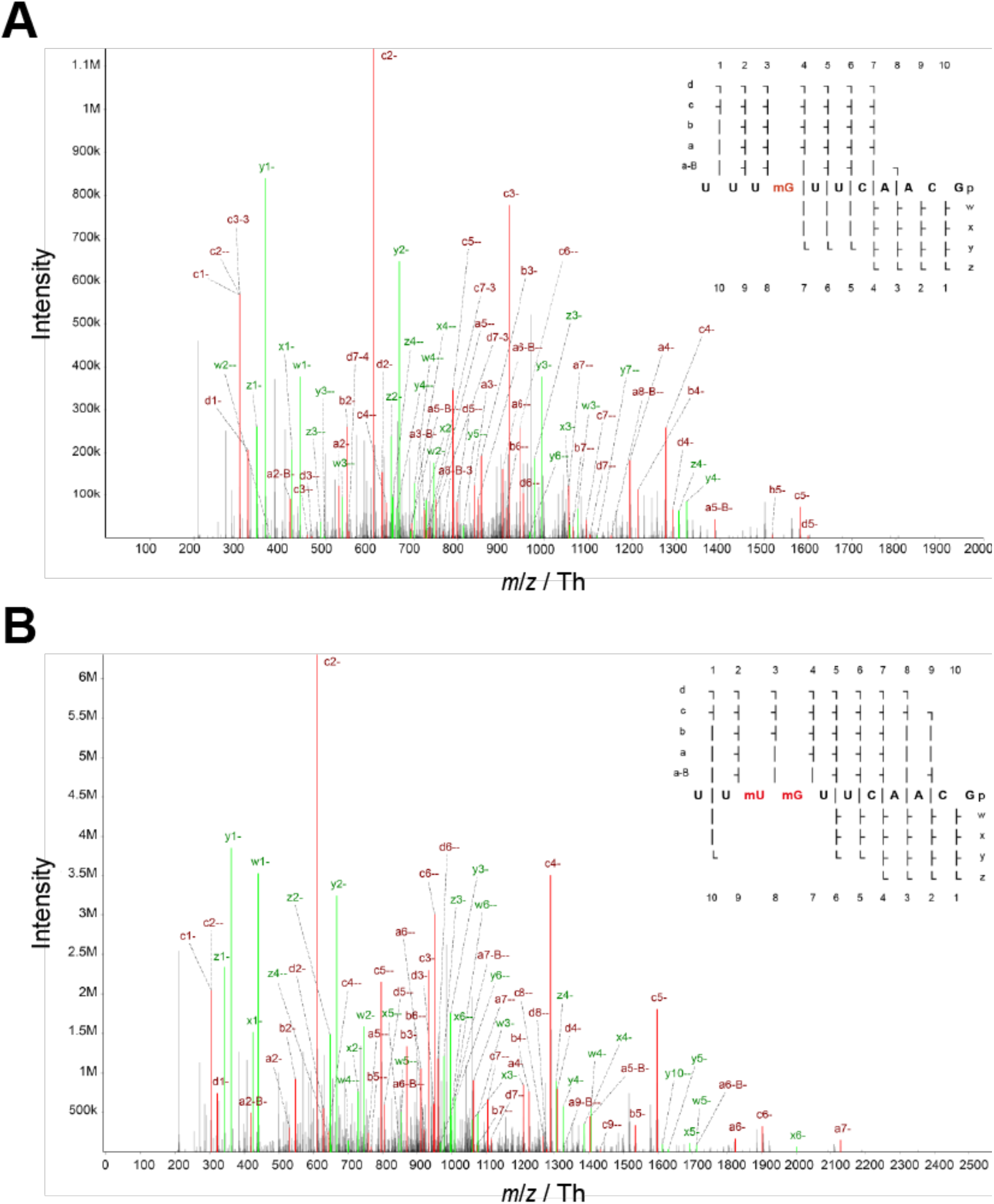
Mass spectrometric characterisation of 16S mt-rRNA oligonucleotides. Annotated RNA MS^2^ fragmentation spectra from RNase T1 cleavage products of (A) U1369/Gm1370 (maximum ion score: 198.87; Q value: 0) and (B) Um1369/Gm1370 (maximum ion score: 220.06; Q value: 0) modified 16S mt-rRNA. Scores were generated during assignment by NucleicAcidSearchEngine. Related to Figure 6B.

**Figure S11.**
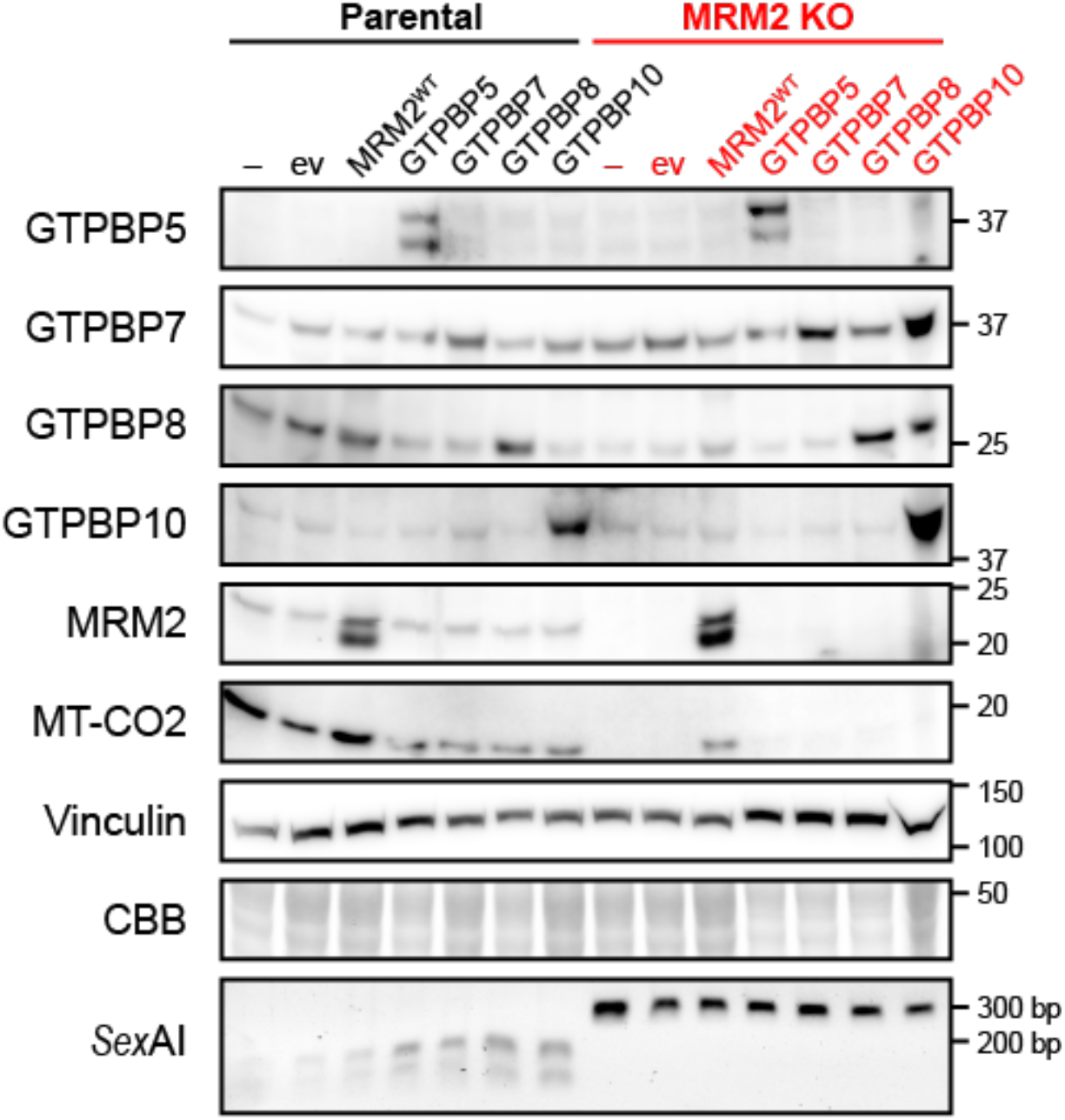
Investigation on the functional rescue of mitochondrial translation in MRM2-depleted cells by mitochondrial GTPBPs. Immunoblot evaluation of functional rescue of mitochondrial translation in cell lines complemented with wild-type MRM2 (MRM2^WT^), as well as GTPBP5/MTG2, GTPBP7/MTG1, GTPBP8 and GTPBP10. Molecular weights of protein standards are presented in kDa to the right of each blot. Coomassie brilliant blue (CBB) staining is shown as a loading indicator. For each cell line, electrophoretically separated *Sex*AI-digested amplicons of the genomic *MRM2* locus targeted for gene editing are presented.

**Table S1.** CryoEM data collection parameters.

|  | **Non-tilted**<br>**data collection** | **Tilted**<br>**data collection** |
| --- | --- | --- |
| **Data collection and processing** |  |  |
| Microscope | Titan Krios | Titan Krios |
| Detector | Falcon 3EC | Falcon 3EC |
| Magnification | 75,000x | 75,000x |
| Voltage (kV) | 300 | 300 |
| Electron exposure (e<sup>-</sup> Å<sup>-2</sup>) | 52.5 | 52.1 |
| Defocus range (μm) | -2.6 to -1.0 | -2.2 to -1.4 |
| Pixel size (Å) | 1.06 | 1.06 |
| Stage tilt (°) | 0 | -20 |
| Initial particle images (no.) | 1,014,736 | 1,694,785 |
| Final particle images (no.) | 510,450 | 681,420 |

**Table S2.**
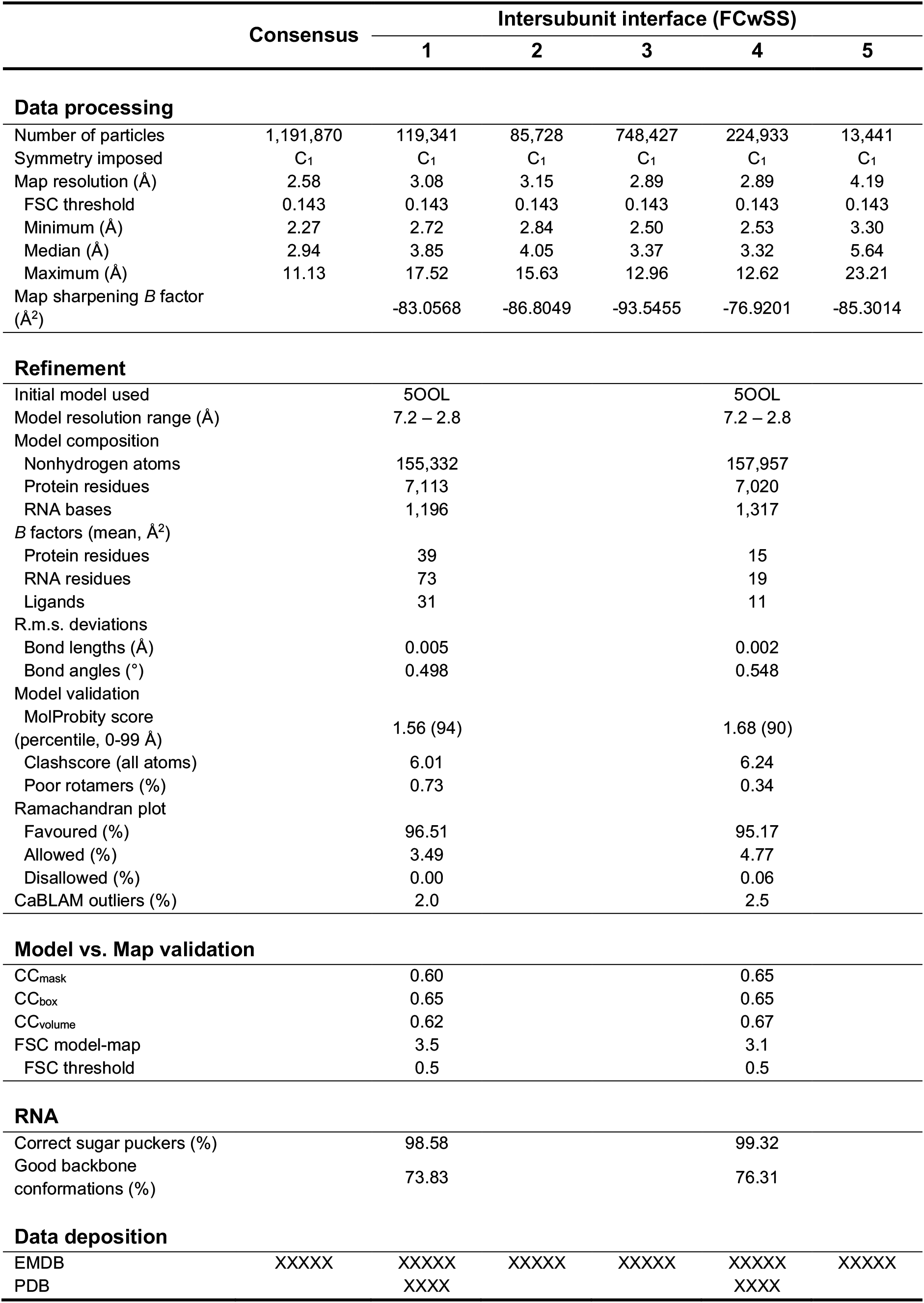
CryoEM map refinement and model validation statistics – consensus and intersubunit interface focused classification with signal subtraction.

**Table S3.**
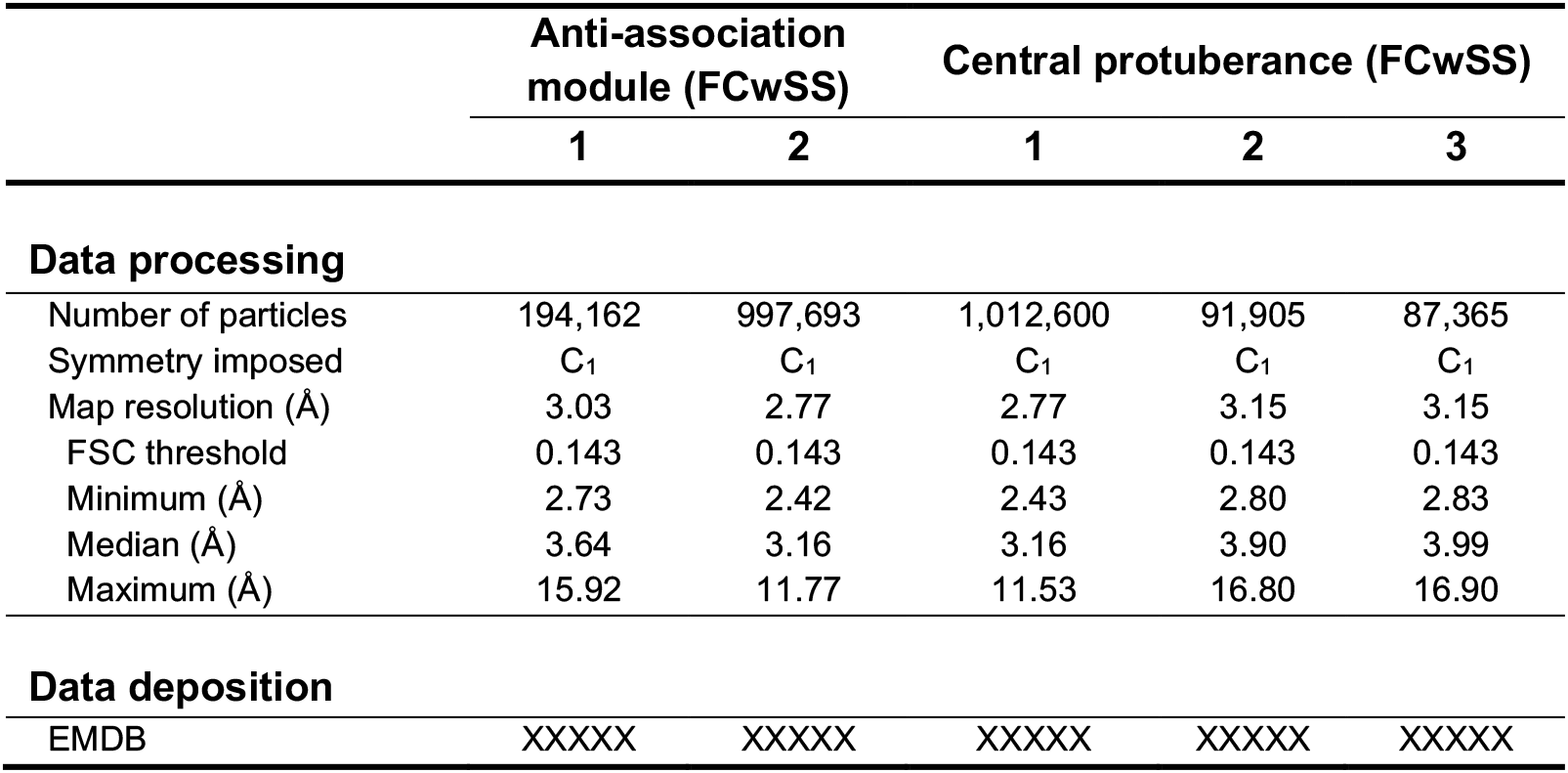
CryoEM map refinement – anti-association module and central protuberance focused classification with signal subtraction.

**Movie S1. Inspection of structural heterogeneity in mtLSU particles in the absence of MRM2.**

UMAP visualisation of the latent space is presented on the right, with the on-data cluster centres represented as diamonds. The latent space exploration path though cluster centres (in no particular order) can be followed by a red dot, which corresponds to the volume (latent space representation) shown on the left. All volumes are presented with the same isosurface threshold.

**Data S1. The code for calculating RT-stop scores and cleavage protection scores is found in this paper’s supplemental information.**

## RESOURCE AVAILABILITY

### Lead contact

Further information and requests for resources and reagents should be directed to and will be fulfilled by the lead contact, Michal Minczuk.

### Materials availability

Materials generated in this study are available upon request.

### Data and code availability

Sequencing datasets generated in this manuscript have been deposited at GEO and are publicly available as of the date of publication. Accession numbers are listed in the key resources table. The cryoEM density maps including the 2.58 Å-resolution map of the consensus mtLSU from *MRM2* knock-out cells, all maps obtained by focused classification with signal subtraction (FCwSS) and all structural models have been deposited in EMDB and PDB, and are publicly available as of the date of publication. Accession numbers are listed in the key resources table.

Custom R scripts used for RNA mass spectrometry analysis are property of Storm Therapeutics. All other original code is available in this paper’s supplemental information.

Any additional information required to reanalyse the data reported in this paper is available from the lead contact upon request.

## EXPERIMENTAL MODEL AND SUBJECT DETAILS

Unless stated otherwise, HEK 293T Flp-In T-REx and HEK 293T were maintained in high glucose DMEM with GlutaMAX and sodium pyruvate supplemented with 10% FBS, 1x penicillin-streptomycin. All cells were kept at 37 °C and in a 5% CO2 atmosphere. Stocks used in this study are reported in the key resources table.

*Drosophila melanogaster* flies were maintained in a humidified temperature-controlled incubator set to 25 °C with 12 h/12 h light/dark cycles. Crosses and respective progenies were raised in a humidified incubator at 29 °C. Food consisted of agar, cornmeal, molasses, propionic acid and yeast. Stocks used in this study are reported in the key resources table.

## METHOD DETAILS

### Knock-out cell line generation

HEK 293T Flp-In T-REx cells were transfected with plasmids encoding the (+) and (-) ZFNs target to *MRM2* using the Cell Line Nucleofector Kit V with the A-23 program in the Nucleofector 2b device, according to the supplier’s recommendations. After recovery, clonal lines were screened by immunodetection of MRM2 and subsequent DNA sequencing (amplification with MRM2 Fw and MRM2 Rv). The generated knock-out cells were cultured in medium supplemented with 20% FBS and 50 mg mL^-1^ uridine.

### RNA extraction for library preparation

RNA was extracted from parental and HEK 293T Flp-In T-REx cells in which *MRM1*, *MRM2* or *MRM3* was knocked-out. For each condition 3 biological replicates were obtained (one 15 cm plate each). Mitochondria isolation and RNA extraction was conducted as previously described (Sas-Chen, Nir and Schwartz, 2021). In brief, collected cells were resuspended in mannitol-containing mitochondria extraction buffer and incubated on ice for 20 minutes. Cells were then homogenized by running the suspension 15 times through a 25G needle, followed by a series of low- and high-speed centrifugations. The resulting pellet, representing the mitochondria-enriched fraction, was used as input for RNA purification using TRI Reagent according to the manufacturer’s instructions.

### Library preparation and sequencing

Strand-specific libraries were generated on the basis of previously described protocols (Engreitz et al., 2013; Shishkin et al., 2015; Marchand et al., 2016; Incarnato et al., 2017; Sas-Chen, Nir and Schwartz, 2021). In brief, for each sample, 200 ng of mitochondria-enriched RNA was fragmented by mixing the RNA in bicarbonate buffer (pH 9.2) and heating for 5 minutes at 92 °C. Next, samples were subjected to FastAP Thermosensitive Alkaline Phosphatase and Turbo DNase treatments, followed by a 3′ ligation of RNA adaptor 1 using T4 ligase. Ligated RNA was separated into two identical pools and reverse transcribed using SuperScript III Reverse Transcriptase in two distinct concentrations of deoxyribonucleotide triphosphates (dNTPs; final concentration 2 μM and 500 μM), using the RT primer. As the used RT-enzyme stalls at 2’-*O*-methlated sites in low dNTP conditions, the first pool, subjected to 2 μM dNTPs was later used to obtain RT-stop signals (Incarnato et al., 2017). The second pool, transcribed using 500 μM dNTPs, was used to obtain the background signal for RT-stops (elaborated in “Identification of putative 2’-*O*-methylation sites”), as well as to calculate cleavage protection scores (‘score A’; Marchand et al., 2016). Resulting cDNA was subjected to a 3′ ligation with adaptor 2 using T4 ligase. The single-stranded cDNA product was then PCR amplified for 9-12 cycles. Libraries were sequenced on a NextSeq 500 platform generating short paired-end reads, ranging from 25 to 55 bp from each end.

### Identification of putative 2’-*O*-methylation sites

A reference genome for alignment of reads was generated on the basis of chrM from the GENCODE release 32 (GRCh38.p13). Reads were mapped using the STAR aligner (Dobin et al., 2013) with *–alignIntronMax* set to 1. To identify methylated sites, coverage and number of reads starting and ending at each position was counted using bam2ReadEnds.R (García-Campos, 2019; Garcia-Campos et al., 2019).

Determination of RT-stop scores. Under limiting dNTP concentrations, the RT stops one nucleotide downstream of the methylated site. Thus, for each position, an RT-stop ratio was calculated by dividing the number of reads stopping at the downstream position by the coverage at that position. RT-stop ratio of low dNTP samples was then divided by the RT-stop ratio of the high dNTP sample, which resulted in a fold change RT-stop score representing enrichment of the 2’-*O*-methylated site in low dNTP versus the background signal. Significance of changes in methylation level in wild-type and knock-down cells was assessed using Student’s t-test comparing the calculated fold change RT-stop scores in the 3 biological replicates of each condition.

Determination of cleavage protection scores. Cleavage protection scores were determined as in the calculation of ‘score A’ in (Birkedal et al., 2015). The length of the flanking region used for calculation was set to 6 nucleotides, centred on the measured site. Scores were calculated only for positions with a median of >15 reads per position in the flanking 12 nucleotides surrounding the measured site. Significance of changes in methylation level in parental and knock-out cells was assessed using Student’s t-test comparing the calculated ‘score A’ in 3 biological replicates of each condition. The code for calculating RT-stop scores and cleavage protection scores is found in this paper’s supplemental information (**Data S1**).

### Monitoring of cell proliferation

Cells were plated in glucose-free DMEM supplemented with 10% FBS, 1x GlutaMAX, 1 mM sodium pyruvate, 1x penicillin-streptomycin and either 4.5 g L^-1^ glucose or 0.9 g L^-1^ galactose. Cell confluence was measured every 4 hours using the Incucyte ZOOM in High Definition Imaging Mode and analysed with the associated software.

### Assessment of mitochondrial respiration

Cells were seeded and allowed to adhere for at least 6 h to Seahorse XF96 cell culture microplates previously coated with poly-L-lysine. Culture medium was removed and the cell layer washed with assay medium (DMEM without sodium bicarbonate, 4.5 g L^-1^ glucose, 1x GlutaMAX, 1 mM sodium pyruvate, 150 mM NaCl, pH 7.4). After equilibration in assay medium (1 h, 37 °C) with atmospheric CO2, oxygen consumption rate (OCR) and extracellular acidification rate (ECAR) were measured in a Seahorse XF96 Extracellular Flux Analyzer, using sequential injections of 2 μM oligomycin, 0.35 μM BAM15, and 1 μM rotenone and 1 μM antimycin A. OCR and ECAR values were normalised by the total amount of DNA in each well, determined using the CyQUANT Cell Proliferation Assay kit.

### Spectrophotometric assessment of MRC activity

The spectrophotometric measurement of the activity of MRC complexes was performed with small modifications to the protocol described in (Kirby et al., 2007). Measured activities are normalised to the activity of citrate synthase, and values are presented relative to those of the parental cell line.

### PAGE and immunodetection of proteins

Proteins were denatured, separated by SDS-PAGE in Bolt 4-12% Bis-Tris Plus gels, and transferred onto nitrocellulose membranes via dry transfer. These were blocked with 5% non-fat milk in PBST for 1 h at room temperature, and then incubated overnight with primary antibodies at 4 °C. After three washes, membranes were incubated with the appropriate HRP-conjugated secondary antibody for 1 h at room temperature, washed and developed with ECL reagent using an Imager 680.

BN-PAGE was performed on whole cells following the protocol by (Fernandez-Vizarra and Zeviani, 2021).

Primary and secondary antibodies are listed in the key resources table.

### In-gel assessment of MRC activity

Following the resolution of whole cell lysates by BN-PAGE, gels were incubated with the following solutions at room temperature: Complex I IGA solution (5 mM Tris-HCl pH 7.4, 1 mg mL^-1^ nitro blue tetrazolium, 0.1 mg mL^-1^ NADH), Complex II IGA solution (5 mM Tris-HCl pH 7.4, 1 mg mL^-1^ nitro blue tetrazolium, 0.2 mM phenazine methosulfate, 20 mM succinate), Complex IV IGA solution (50 mM potassium phosphate buffer pH 7.4, 1 mg mL^-1^ 3,3’-diaminobenzidine tetrahydrochloride hydrate, 1 mg mL^-1^ cytochrome c, 24 U mL^-1^ catalase, 75 mg mL^-1^ sucrose). Upon appearance of coloured bands, gels were washed with water and scanned under white light.

### Labelling of *de novo* synthesised proteins

Cells of interest were incubated twice in methionine/cysteine-free DMEM for 10 min, and then in methionine/cysteine-free DMEM supplemented with 5% dialysed FBS, 1x GlutaMAX, 1 mM sodium pyruvate, 96 μg mL^-1^ L-cysteine for 20 min, after which 100 μg mL^-1^ emetine were added. After 30 min, 100 μCi mL^-1^ of [^35^S]-L-methionine were added and labelling proceeded at 37 °C for 1 h. Cells were collected and washed with PBS before being lysed by addition of 0.1% DDM, 1.7 U μL^-1^ benzonase and 1x cOmplete protease inhibitor cocktail. Sample components were separated in 10-20% Tris-glycine SDS-PAGE gels, which were stained with Coomassie Brilliant Blue before being dried. The autoradiogram was obtained by exposing the dried gel to a phosphor screen and developed using a Typhoon Biomolecular Imager. Densitometry profiles and baseline-corrected quantification were performed using ImageJ (Schindelin et al., 2012).

### Isolation of mitochondria

Cells were detached from culture vessels and incubated on ice in hypotonic buffer (20 mM HEPES pH 7.8, 5 mM KCl, 1.5 mM MgCl2, 2 mM DTT, 1 mg mL^-1^ BSA, 1 mM PMSF, 1x cOmplete protease inhibitor cocktail) for 10 min. A Balch-type homogeniser was used to mechanically lyse cells by 5 passes through the chamber loaded with a 12 μm clearance ball bearing. After homogenisation, 2.5x MSH buffer (50 mM HEPES pH 8.0, 525 mM mannitol, 175 mM sucrose, 5 mM EDTA, 5 mM DTT, 2.5 mg mL^-1^ BSA, 2.5 mM PMSF, 2.5x cOmplete protease inhibitor cocktail) was added. Debris were pelleted by centrifugation at 1,000 *g* for 10 min, and crude mitochondria subsequently pelleted from the supernatant at 10,000 *g* for 20 min. These were layered on top of a discontinuous density gradient (1.5 M, 1.0 M, and 0.5 M sucrose in 10 mM HEPES pH 7.8, 5 mM EDTA) and ultracentrifuged at 120,000 *g* for 1 h. The mitochondrial layer was washed in 1x MSH and pelleted at 10,000 *g*. All manipulations and centrifugations were performed at 4 °C.

### Assessment of mitoribosome composition

The procedure followed that described for quantitative analysis of density gradients by mass spectrometry (qDGMS) in (Páleníková et al., 2021). In brief, cells were cultured in DMEM for SILAC, supplemented with 20% dialysed FBS, 50 mg mL^-1^ uridine, 1x penicillin-streptomycin, 200 mg L^-1^ L-proline, and either 0.398 mM L-arginine and 0.798 mM L-lysine, or 0.398 mM ^13^C6,^15^N4-L-arginine and 0.798 mM ^13^C6,^15^N2-L-lysine. Cell populations were allowed to expand separately in these media for at least 7 doubling periods, with frequent media exchange. Upon harvesting, cell populations were mixed 1:1 based on total protein content. Isolated mitochondria were lysed (50 mM Tris-HCl pH 7.4, 150 mM NaCl, 1 mM EDTA, 1% Triton X-100, 1x cOmplete EDTA-free protease inhibitor cocktail, 2 U μL^-1^ SUPERase•In RNase inhibitor), clarified and loaded on a continuous 10-30% (w/v) sucrose density gradient (50 mM Tris-HCl pH 7.4, 100 mM NaCl, 20 mM MgCl2, 1x cOmplete EDTA-free protease inhibitor cocktail) formed with a Gradient Station. Gradients were spun at 100,000 *g* in a TLS-55 swinging bucket rotor for 135 min at 4 °C. Fractions were manually collected from the top of each gradient. Results for each mix were corrected for imperfect mixing by taking into consideration the median ratio of labels from mass spectrometric analysis of whole cell mix lysates.

### Protein mass spectrometry

Liquid samples were precipitated with 20 vol. of cold ethanol and incubated at -20 °C for 16 h. Precipitates were collected by centrifugation, redissolved in 50 mM ammonium bicarbonate buffer (pH 8.0) and digested with trypsin (12.5 ng µL^-1^, 37°C, overnight). Peptide mixtures were resolved by reverse-phase UPLC on an EASY-nanoLC1000 and an Acclaim PepMap C18 column (50 µm x 150 mm, 2 μm, 100 Å, 300 nL min^-1^) using an 84 min gradient of 5% to 40% acetonitrile with 0.1% formic acid, followed by an increase in acetonitrile concentration to 90% and re-equilibration with 5% acetonitrile, within 105 min. Peptides were analysed by positive ion electrospray mass spectrometry using a Q Exactive plus mass spectrometer and a routine to fragment and analyse the 10 most abundant multiply charged peptide ions each second. Full scan MS data (400 to 1600 *m/z*) were recorded at a resolution of 70,000 with an automatic gain control (AGC) target of 10^6^ ions and a maximum ion transfer of 20 ms. Ions selected for MS^2^ were analysed using the following parameters: resolution 17,500; AGC target of 5 x 10^4^; maximum ion transfer of 100 ms; 2 *m/z* isolation window; for HCD, a normalized collision energy 28% was used; and dynamic exclusion of 20 s. A lock mass ion (polysiloxane, *m/z* = 445.1200) was used for internal MS calibration. Proteins were identified using the MaxQuant software package. Peptide information was filtered using Perseus, removing proteins only identified by site that matched a decoy database of random peptides and contaminants. Preliminary assessment of incorporation of heavy amino acids was performed by processing the peptide information of the heavy-only labelled samples using MaxQuant.

### Mitochondrial ribosome footprinting

Cells were incubated with 200 μg mL^-1^ chloramphenicol for 10 min and 200 μg mL^-1^ cycloheximide for an additional 5 min. After washing with PBS, the monolayer was flash-frozen on liquid nitrogen and thawed in lysis buffer (20 mM Tris-HCl pH 7.5, 150 mM NaCl, 5 mM MgCl2, 1 mM DTT, 1% Triton X-100, 200 μg mL^-1^ chloramphenicol, 200 μg mL^-1^ cycloheximide). Half of the clarified lysates was incubated with 375 mU μL^-1^ RNAse I at 28 °C for 30 min, and monosomes recovered by ultracentrifugation in a continuous 10-30% (w/v) sucrose density gradient containing chloramphenicol and cycloheximide. After 40 mU μL^-1^ proteinase K incubation at 42 °C for 30 min, ribosome-protected fragments were extracted in acid phenol-chloroform. Total RNA was extracted from the other half using TRIzol LS, DNA degraded using TURBO DNase and mt-rRNA depleted using the Ribo-Zero Gold kit before alkaline fragmentation (45 mM sodium bicarbonate, 5 mM sodium carbonate, 1 mM EDTA, 95 °C, 15 min). Both RNA sample sets were subjected to size selection (25 nt to 35 nt) after TBE-Urea PAGE. Following, RNA was extracted from gel pieces (300 mM sodium acetate pH 5.5, 0.25% SDS, 4 °C, 18 h) and standard library preparation was carried out using the TruSeq Small RNA Library Preparation kit (Illumina). Libraries were sequenced (50 bp single read) in the CRUK Cambridge Institute Genomics Core using a HiSeq 4000 system. Adaptor sequences were trimmed from reads using FASTX-Toolkit (http://hannonlab.cshl.edu/fastx_toolkit/), trimmed reads mapping to nuclear-encoded ribosomal RNA (rRNA) were discarded, and the remaining reads were mapped sequentially to mitochondrial rRNA (mt-rRNA); mitochondrial transfer RNA (mt-tRNA); and mitochondrial messenger RNA (mt-mRNA) using bowtie version 1 (Langmead et al., 2009), with parameters *-v 2 --best*. To normalise for library size, read counts per mitochondrial gene were expressed as reads per million (RPM) mapping in the positive-sense orientation to nuclear-encoded mRNA (from NCBI RefSeq) in each library. When calculating mitochondrial gene ribosomal occupancy and translation efficiency (i.e. RiboSeq RPM / RNASeq RPM), overlapping CDS regions in ATP8/ATP6 and ND4L/ND4 were excluded due to ambiguous assignment. Additionally, reads with 5’ ends mapping within 15 nt of the start codon or 45 nt of the stop codon of each gene were excluded to avoid counting ribosomes paused during initiation or termination of translation.

### Purification of mtLSU

Mitochondria were purified from *MRM2* knock-out cells as described in “Isolation of mitochondria”, and lysed in 20 mM HEPES, 300 mM KCl, 50 mM MgCl2, 1 mM DTT and 1% Triton X-100 supplemented with RNase and EDTA-free protease inhibitors.

The clarified lysate was loaded on a continuous 10-30% (w/v) sucrose density gradient (20 mM HEPES pH 7.6, 300 mM KCl, 5 mM MgCl2, 1 mM DTT) and ultracentrifuged at 100,000 *g* for 135 min. Fractions enriched in mtLSU were pooled and ultracentrifuged at 625,700 *g* for 35 min. The obtained pellet was resuspended in lysis buffer without Triton X-100 and loaded on a second continuous 10-30% (w/v) sucrose density gradient, which was ultracentrifuged at 100,000 *g* for 12 h. Fractions were automatically collected using an ÄKTAprime plus system with a 60% (w/v) sucrose chase solution. The absorbance profile at 260 nm was monitored and fractions enriched in mtLSU were pooled. Ribosomal particles were pelleted at 418,000 *g* for 2 h. The pellet was washed in buffer without sucrose or DTT, resuspended, briefly centrifuged to remove aggregates, and used for cryoEM grid preparation. All manipulations and centrifugations were performed at 4 °C.

### CryoEM grid preparation and data collection

Quantifoil holey carbon R2/2 mesh 300 copper grids were glow discharged at 20 mS for 60 s. Sample grids were prepared using a Vitrobot Mark IV set to 95% relative humidity and 4 °C. Purified mtLSU (∼0.2 μg μL^-1^ total protein) was applied to the carbon-coated side of grids (3 μL), blotted (15 s waiting time, 2.5 s blotting time, 0 blotting force) and plunge frozen in liquid ethane.

Data were collected at the cryoEM facility of the Biochemistry Department (University of Cambridge) on a 300 keV Titan Krios equipped with a Falcon 3EC direct electron detector in integrating mode. Due to preferential orientation of the particles, a second dataset was collected where the grid was tilted by 20°, as suggested by cryoEF (Naydenova and Russo, 2017) using data from the processed non-tilted dataset. A total of 10,078 and 10,401 micrographs were acquired for the non-tilted and tilted datasets, respectively. In both cases a total dose of ∼52 e^−^ Å^-2^ was distributed over 18 frames, with an exposure time of 600 ms; C2 and objective apertures were 70 μm and 100 μm, respectively (**Table S1**).

### CryoEM data processing and model building

Acquired data was processed (**Figure S5**) using RELION-3.1 (Zivanov et al., 2018; Zivanov, Nakane and Scheres, 2020). Beam-induced motion correction was performed using MotionCor2 (Zheng et al., 2017) and CTF estimation using CTFFIND-4.1 (Rohou and Grigorieff, 2015). Particles were picked using the Laplacian of Gaussian (LoG) filter and imported into cryoSPARC (Punjani et al., 2017) where an initial reference was generated *ab initio* and used for clearing the stack by 3D heterogeneous refinement. Clean particle stacks were processed (3D auto-refinement, CTF refinement, Bayesian polishing) individually with RELION-3.1 and then used for 3D classification. Particles from both datasets that led to reconstruction of maps with detailed features were merged to generate a partial consensus map (846,646 particles); all particles were used to generate a full consensus (1,191,870 particles). Focused 3D classifications (**Figure S7**, **Figure S9**, **Table S2** and **Table S3**) were performed using RELION-3.1 by subtracting signal outside a mask over the regions of interest (FCwSS) and classifying particles without updating alignments; classes of interest were further 3D auto-refined. Masks used for FCwSS are presented in **Figure S7** and **Figure S9**.

Full models of states 1 and 4 (**Figure 5**) were built and refined. The structure of a previously reported mitochondrial assembly intermediate (PDB: 5OOL) was used as reference model. This model was first fit into the density using Chimera (Pettersen et al., 2004). Iterative cycles of model building and refinement of ribosomal proteins and rRNA were performed using *Coot* (Emsley et al., 2010) and PHENIX real-space refinement (Adams et al., 2010), respectively. The model was validated with MolProbity (Chen et al., 2010) and the PHENIX suite (Adams et al., 2010). Model building statistics are presented in **Table S2**.

Heterogeneity analysis was performed using cryoDRGN (Zhong et al., 2021). A 1,191,870 particles stack, downsampled to 256 px (1.49 Å px^-1^) was used, alongside poses and CTF parameters extracted from the reconstructed full consensus (**Figure 5A** and **Figure S5**), to train the variational auto-encoder for 25 epochs with a 10D latent variable and a 1024x3 architecture. The distribution of latent encodings was visualised with the built-in 2D uniform manifold approximation and projection (UMAP) and *k*-means clustering was performed with *k* = 30. A trajectory connecting the cluster centres in latent space was determined and volumes of anchor points along that path generated “on-data” by evaluating the trained decoder with their latent variable values. Additional cluster centres were determined by selecting a region of interest in the UMAP representation using the cryoDRGN-generated Jupyter Notebook, determining the median coordinates for the selected datapoints and identifying the closest datapoint that represents a particle from the dataset.

Figures of maps and models were generated using ChimeraX (Pettersen et al., 2021).

### Lentivirus production and stable cell line generation

Lentivirus were produced in HEK 293T transiently co-transfected with pWPXLD:IRES:PuroR containing the transgene of interest, psPAX2 and pMD2.G using FuGENE 6 transfection reagent according to the manufacturer’s instructions, using a DNA mass to transfection reagent volume ratio of 0.33. Lentiviral supernatants collected after 48 h were spun, filtered and split equally among cells of interest. Cells that successfully integrated the transfer plasmid were selected with 1 μg mL^-1^ puromycin.

To verify the genetic background of cell lines, the genomic region modified to knock-out *MRM2* was amplified and digested with *Sex*AI. Reaction products were separated by electrophoresis in 1% agarose TBE gels.

### RNA oligonucleotide mass spectrometry

Large mitoribosomal subunits were purified from cultured cells using a continuous sucrose density gradient as described in “Purification of mtLSU”. Subsequently, RNA was extracted from selected fractions using TRIzol LS and screened for purity and concentration using a 2200 TapeStation system. Oligonucleotides were prepared from RNA using RNAse T1 and chromatographically separated by ion pair reverse phase chromatography (200mM HFIP, 8.5 mM TEA in water as eluent A, and 100 mM HFIP, 4.25 mM TEA in methanol as eluent B). The oligonucleotides were resolved by a non-linear gradient of 2.5% to 20% B at 200 nL min^-1^ on Acclaim PepMap C18 solid phase and characterised by negative ion tandem LC-MS in a Q Exactive HF hybrid quadrupole-orbitrap. Data were collected in data-dependent acquisition mode, with full scan MS^1^ data acquired between 700 and 3500 *m*/*z*. The top five ions generating signals with the most intense signals were selected for fragmentation and subsequent MS^2^ characterisation. Tandem MS data was analysed using the OpenMS software suite (Röst et al., 2016), and custom R scripts. Oligonucleotide database searching was performed using NucleicAcidSearchEngine, according to parameters given in (Wein et al., 2020). Briefly, isotopologues between the monoisotopic and +4 peak were considered for assignment with an accuracy of 3 ppm at MS^1^ and MS^2^, and single sodium and potassium ions were considered as adducts, in addition to single methylation of each nucleobase as a variable modification. Label-free quantification of oligonucleotide precursors was initially assessed using an updated version of FeatureFinderID (Weisser and Choudhary, 2017) and confirmed by manual MS^1^ peak integration.

### Quantification of transcript levels

Levels of *CG11447* (*DmMRM2*) and *αTub84B* transcripts were determined by RT-qPCR. Total RNA was extracted from 3 larvae using TRIzol reagent, genomic DNA removed with TURBO DNA-free Kit, and reverse transcribed with Maxima H Minus cDNA Synthesis Master Mix. The reaction products were amplified with PowerUp SYBR Green Master Mix using primer pairs targeting *DmMRM2* (DmMRM2 Fw and DmMRM2 Rv) and *αTub84B* (αTub84B Fw and αTub84B Rv). qPCR was performed in a QuantStudio 3 Real-Time PCR System. No-RT and no-template control reactions were included. Specificity of primer pairs was evaluated by the melting profile, and the amplification efficiency was 1.93 and 1.95 for the pair targeting *DmMRM2* and *αTub84B*, respectively. Transcript levels were determined using the comparative Ct method with amplification efficiency correction and normalised for the average of *αTub84B* (Pfaffl, 2001).

### Locomotor assay

The startle-induced negative geotaxis (climbing) assay was performed using a counter-current apparatus. Briefly, adult male flies were placed into the first chamber, tapped to the bottom, and given 10 s to climb a 10 cm distance. This procedure was repeated five times (five chambers), and the number of flies that has remained into each chamber counted. The weighted performance was normalized for each genotype to the maximum possible score and expressed as climbing index (Greene et al., 2003).

## QUANTIFICATION AND STATISTICAL ANALYSIS

Individual experimental values with average, mean ± SD or median values are presented. Statistical parameters, tests and significance, as well as number of replicates (*n*) can be found accompanying the respective data in figure legends.

## SUPPLEMENTAL INFORMATION

Supplemental information includes eleven figures, three tables, one movie and one R script for calculation of ribose methylation scores. These can be found online at https://www.doi.org/XXXX.

## ACKNOWLEDGEMENTS

The authors would like to acknowledge Dima Chirgadze (cryoEM facility, University of Cambridge) for assistance with grid screening and acquisition of electron microscopy data, Michael Harbour, Shujing Ding and Ian M. Fearnley (Mass Spectrometry facility, MRC MBU, University of Cambridge) for their help in the proteomics analysis, Merlin Hartley and Andrew Raine for IT support, and the members of the MRC MBU Mitochondrial Genetics group and Daniel Grba for insightful discussions.

This work was supported by core funding from Medical Research Council UK (MC_UU_00015/4, MC_UU_00015/6), Fundação para a Ciência e a Tecnologia (PD/BD/105750/2014) to P.R.-G., a specialist Programme from Blood Cancer UK (12048), the Kay Kendall Leukaemia Fund, the UK MRC (MR/T012412/1), a Wellcome Trust strategic award to the Cambridge Institute for Medical Research (100140), a core support grant from the Wellcome Trust and MRC to the Wellcome Trust – MRC Cambridge Stem Cell Institute, the Cambridge National Institute for Health Research Biomedical Research Centre and the European Cooperation in Science and Technology (COST) Action CA18233, “European Network for Innovative Diagnosis and treatment of Chronic Neutropenias, EuNet INNOCHRON” to S.P., K.C.D. and A.J. Warren, and European Commission under “Marie Skłodowska-Curie Actions”, Individual Fellowship – Reintegration Panel (Mitobiopath-705560) and Italian Minister of University and Research – Rita Levi Montalcini Program to C.G..

## AUTHOR CONTRIBUTIONS

P.R.-G. and M.M. planned and designed experiments. A.S.-C. and S.S. performed and analysed the transcriptome-wide methylation screening. C.G. performed spectrophotometric assays of MRC activity. A.D., A.E.F. and L.V.H. processed mitoribosome footprinting sequencing. J.F.R. and B.A. performed RNA mass spectrometry. P.R.-G. performed biochemical purifications, K.C.D. collected cryoEM data, P.R.-G. performed image processing and reconstructions with participation of S.P. and K.D, and S.P. built atomic models. L.M.-F. and P.R.-G. performed experiments in the *Drosophila* model. P.R.-G. performed the remaining experiments. P.R.-G. and M.M. drafted the manuscript. M.M., B.A., A.E.F., A.J. Whitworth, A.J. Warren and S.S. supervised the study. All authors revised the manuscript.

## DECLARATION OF INTERESTS

J.F.R. and B.A. are employees of STORM Therapeutics Ltd. M.M. is a founder, shareholder and member of the Scientific Advisory Board of Pretzel Therapeutics, Inc.

